# Macroscopic quantities of collective brain activity during wakefulness and anesthesia

**DOI:** 10.1101/2021.02.03.429578

**Authors:** Adrián Ponce-Alvarez, Lynn Uhrig, Nikolas Deco, Camilo M. Signorelli, Morten L. Kringelbach, Béchir Jarraya, Gustavo Deco

## Abstract

The study of states of arousal is key to understand the principles of consciousness. Yet, how different brain states emerge from the collective activity of brain regions remains unknown. Here, we studied the fMRI brain activity of monkeys during wakefulness and anesthesia-induced loss of consciousness. Using maximum entropy models, we derived collective, macroscopic properties that quantify the system’s capabilities to produce work, to contain information and to transmit it, and that indicate a phase transition from critical awake dynamics to supercritical anesthetized states. Moreover, information-theoretic measures identified those parameters that impacted the most the network dynamics. We found that changes in brain state and in state of consciousness primarily depended on changes in network couplings of insular, cingulate, and parietal cortices. Our findings suggest that the brain state transition underlying the loss of consciousness is predominantly driven by the uncoupling of specific brain regions from the rest of the network.

## Introduction

Interesting phenomena in biological systems are usually collective behaviors emerging from the interactions among many constituents. Large-scale brain activity is not an exception: the brain’s network continuously generates coordinated spontaneous patterns of activity among brain regions at multiple spatiotemporal scales (1-3). Changes in spontaneous brain activity are observed in different brain states, the study of which is essential to understand the organizing principles of brain activity. For instance, anesthesia has been used to transiently induce loss of consciousness and to investigate the neural correlates of the awake state. Previous studies showed that different anesthetics, acting on different molecular targets (4), similarly impact the strength and the structure of functional correlations (5-10), and their dependence on interareal anatomical connections (8, 11). However, how changes in local regions and subnetworks combine to affect the collective brain dynamics and to lose consciousness remains largely unknown. To answer this question, it is essential to precisely characterize the collective properties of different brain states and their dependence on parameters at the system’s level. This dependence is likely not straightforward since, as for many complex systems, the system’s behavior could be differently affected by changes in its parameters. In such a case, while some parameters can largely vary without affecting the system’s behavior (so-called “sloppy” parameters), even small changes in some others can significantly modify it (12-14).

In recent years, statistical mechanics has proven to be more and more useful to describe collective neural activity. Statistical mechanics shows that the behaviors of complex systems can be captured by macroscopic properties, which emerge from the collective activity of the units, in a way largely independent of the microscopic details of the system. These emergent (macroscopic) behaviors can be classified into qualitatively different ordered or disordered phases. Of particular interest are dynamics poised close to phase transitions, or critical points, where order and disorder coexist. Theoretical reasoning shows that complex dynamics and optimal information processing are expected at critical points, making criticality a candidate unifying principle to account for the brain’s inherent complexity necessary to process and represent its environment (15-18). Following this view, studies of whole-brain and local circuits dynamics have proposed that anesthesia shifts the dynamics from the critical point (19, 20). This is supported by the reduction of several measures of brain dynamics complexity under anesthesia (21-24). The global mechanisms underlying different conscious states have been recently investigated using an anatomically-constrained dynamical model with a global coupling parameter in combination with EEG recordings (25). However, it remains unknown which are the macroscopic properties and the relevant local/global parameters describing the transition of collective activity from the awake to anesthetized states. Indeed, different local/global network parameters are likely to jointly determine the different brain states and to differently contribute to the state transitions.

In this study, we addressed these questions by analyzing the brain’s collective activity in different levels of arousal, i.e., during wakefulness and under anesthesia. Specifically, we analyzed resting-state fMRI dynamics of awake and anesthetized macaque monkeys (11). Five different anesthesia protocols, involving 3 different anesthetics (propofol, ketamine, and sevoflurane), were used to induce moderate sedation or deep anesthesia. First, we derived efficient statistics that distinguished between awake and anesthetized brain states. Second, we used these statistics and the maximum entropy principle to model the brain’s activity and to derive important emergent properties that described the different brain states. These emergent properties provided information about the system’s physical state, and about its capability to produce work, to contain information and to transmit it. Finally, we investigated the dependence of collective activity on the different model parameters.

## Results

We analyzed the resting-state fMRI dynamics of five rhesus macaques (*Macaca mulatta*) under different levels of arousal: wakefulness (*n* = 24 scans), two levels of propofol sedation (light, LPP, *n* = 21, and deep, DPP, *n* = 23), sedation through ketamine (KETA, *n* = 22), and two types of sevoflurane anesthesia (SEV2, *n* = 18, and SEV4, *n* = 11) (see Methods and Appendix). fMRI MION time-series were obtained for *N* = 82 previously defined regions of interest (ROIs) (CoCoMac Regional Map parcellation). Each scan was 20 min long and was acquired in time frames of 2.4 s (i.e., 500 time frames).

### Coupling to population reliably distinguished between awake and anesthetized brain states

We were interested on collective patterns displayed among the *N* ROIs, for the six different experimental conditions. For this, we first binarized the z-scored time-series of each ROI, *x_i_*(*t*), by imposing a threshold *θ* = −1 (**Fig. 1A,B**, see Methods). Binarization of time-series has proven to effectively capture and compress fMRI large-scale dynamics (26, 27). We concentrated on different statistics that described the resulting binary data: the activation rate of each ROI, i.e., 〈*σ_i_*〉, the correlation between ROIs, i.e., *C_ij_* = corr(*σ_i_, σ_j_*), and the population activity, i.e., 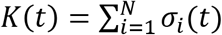 (**Fig. 1C**). We were particularly interested on the coupling of each ROI to the population activity, defined as:

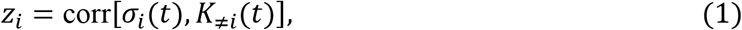

where *K_≠i_*(*t*) is the sum activity of all but ROI *i*: *K_j≠i_*(*t*) = ∑_*j≠i*_ *σ_i_*(*t*). Recent findings showed that propofol anesthesia affects the coupling to global signal in human and rats (28). In the following we showed that the statistics ***z*** = [*z*_1_,…, *z_N_*] provides, with only *N* parameters, a compact description of the binary collective activity and can be used to classify the brain states.

**Figure 1.**
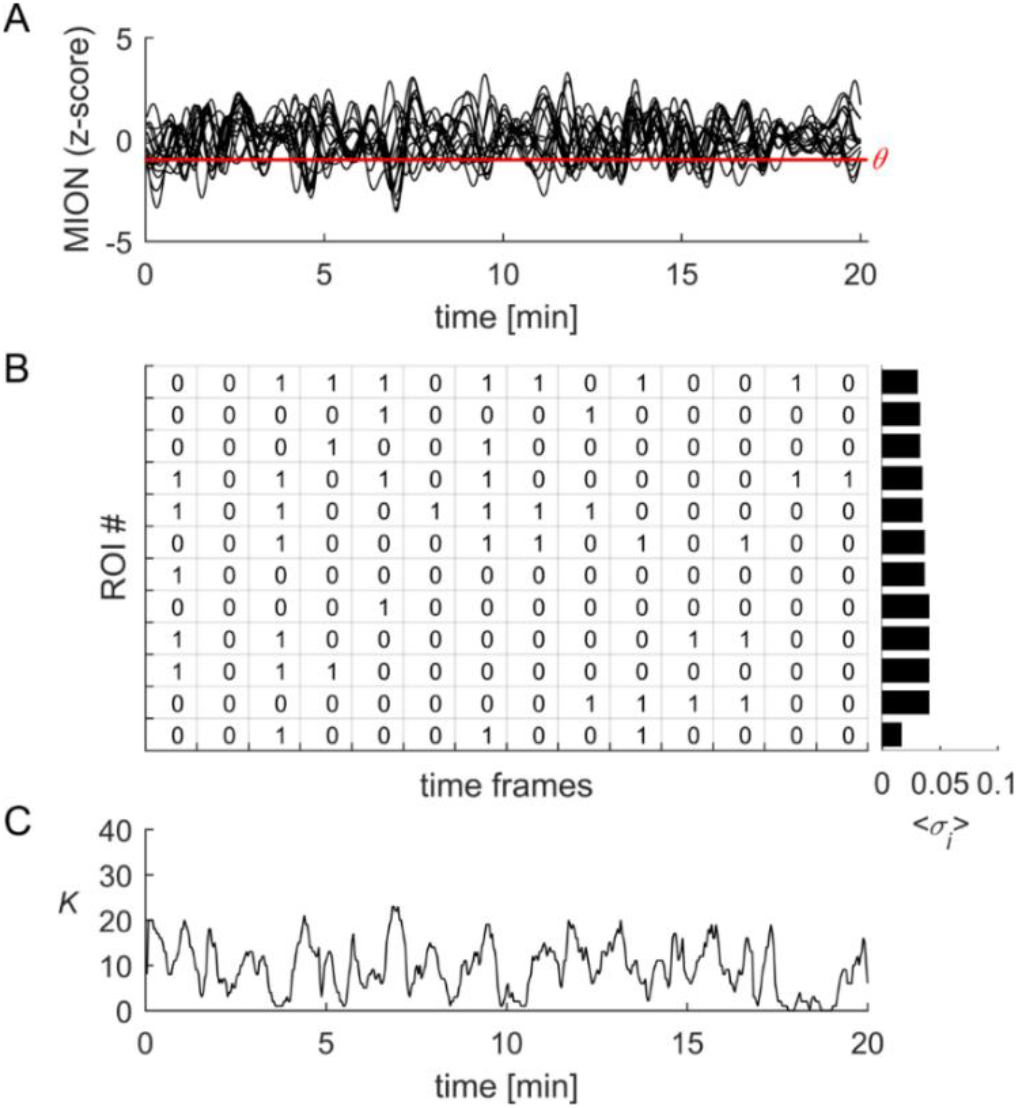
Binarization and statistics. **(A)** MION fMRI signals were z-scored and binarized by imposing a threshold equal to the standard deviation, for each signal. **(B)** In each time bin of 2.4 s, the state of signal of ROI *i*, noted *σ_i_*(*t*), was equal to 1 if the MION signal for this ROI was lower than minus its standard deviation, or equal to 0 otherwise. The average activity of ROI *i* was 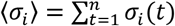, where *n* is the number of time points. **(C)** The population activity was defined as the sum of the binary activity of the *N* ROIs in each time bin *t*, i.e., 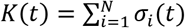.

The couplings to the population were highly predictive of the functional correlations (**Fig. 2A-C**). Indeed, the product *η_ij_ = z_i_ × Z_j_* highly correlated with the functional correlation (FC) between the fMRI time-series of ROIs (*i, j*) (corr.: 0.65–0.78, p < 0.001). Moreover, we found that the vector ***z*** correlated across scans within the same experimental conditions, with the average correlation coefficient being equal to 0.3 ± 0.01 (**Fig. 2D**, *blue* distribution). This correlation was significantly higher (p < 0.001, *F*_(2,3533)_ = 976.5, one-way ANOVA followed by Tukey’s post hoc analysis) than those obtained using the vectors representing the average activities and the correlations, i.e., vectors ***μ*** = [〈*σ*_1_〉,…, 〈*σ_N_*〉 and ***ρ*** = [*C*_1,2_, *C*_1,3_,…, *C*_*N*−1,*N*_], respectively (**Fig. 2D**, *green* and *red* distributions, corr.: 0.06 ± 0.01 and 0.11 ± 0.01).

**Figure 2.**
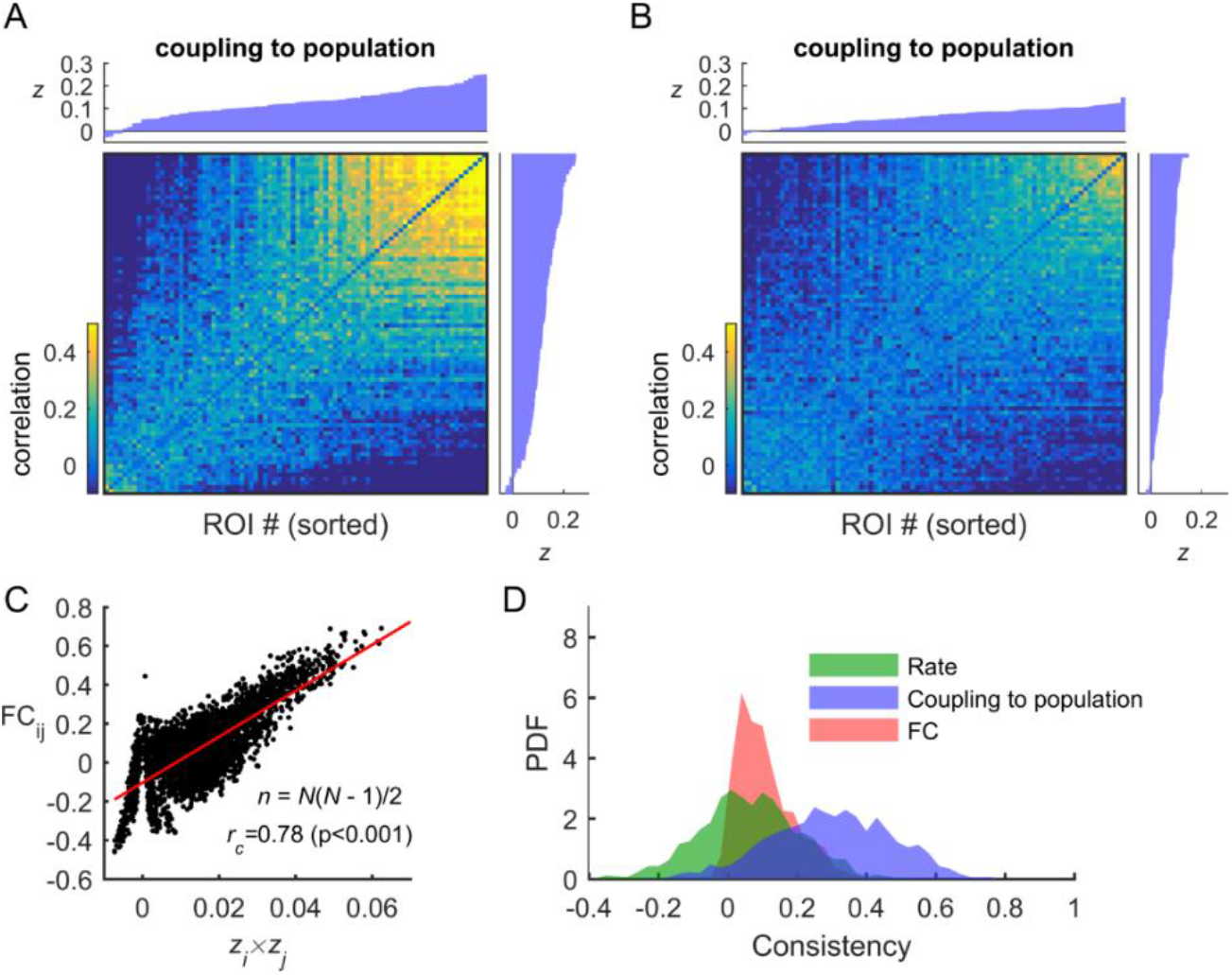
Coupling to population predicts the functional connectivity and is consistent within experimental conditions. **(A)** *Top and left insets:* ROIs were ordered according to *z_i_* averaged for each scan within a given experimental condition (here for the awake condition). *Color map:* The average functional connectivity (FC) is shown after ordering the ROIs according to ***z*. (B)** Same as (A) but for the deep propofol (DPP) anesthesia condition. **(C)**The elements of the FC and the corresponding products of coupling to population (*z_i_z_j_*) highly correlated. **(D)** We tested whether (*σ*), *z*, and FC were similar across scans within the same experimental condition. For example, for the statistic *z*, we calculate the correlation of this *N*-dimensional variable for all pairs of scans belonging to the same experimental condition and computed the distribution of correlation coefficients (*blue* distribution). High correlation coefficients indicate that, within experimental conditions, scans yielded similar vector ***z***. The same can be done for the *N*-dimensional variable 〈*σ*〉 (*green* distribution) and the vector of FC elements (*N*(*N* − 1)/2 dimensions; *red* distribution).

Furthermore, we found that the coupling to the population could be used to classify the awake and anesthesia states with high accuracy (**Fig. 3A**). We tested this by using a classifier based on *k*-means clustering (see Methods). Based on the statistic ***z*** we were able to classify the scans of two categories, awake vs. anesthesia (independently of the anesthetic), with 96.6% of correct classifications (chance level: 50%). This classification performance was higher than the one obtained using the statistics ***μ*** and ***ρ***, yielding 74.0% and 74.8% of correct classifications, respectively (**Fig. 3B,C**). Classification among the six experimental conditions yielded lower performances but was higher for statistic ***z*** than for ***μ*** and ***ρ***: 38.5%, 33.1%, and 29.9%, respectively (chance level: 16.7%). Similar differences in classification performances for population couplings and functional correlations were obtained using continuous (not thresholded) signals (**Fig. S1**). Altogether, these results show that the coupling to population is a reliable marker to distinguish between awake and anesthetized brain states.

**Figure 3.**
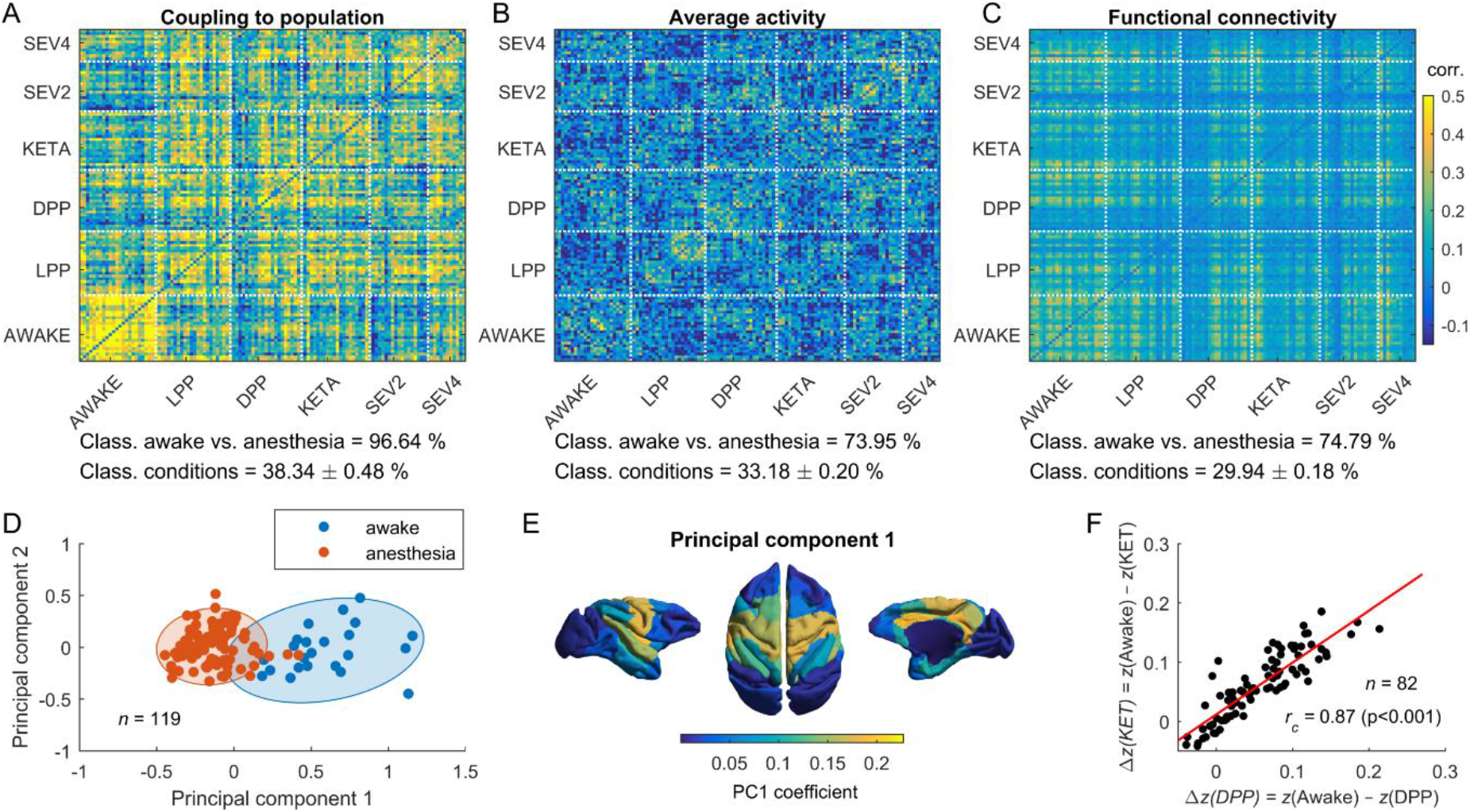
Coupling to population predicts the state of the brain. **(A-C)** Correlation matrix comparing the statistics *z*, 〈*σ*〉, and FC among all scans. For example, in panel (A), the element (*k, l*) of the matrix represents the correlation between the coupling to population vector ***z*** of scans *k* and *l*. Coupling to population clearly separated awake and anesthesia data. Using *k*-means, we evaluated how well the different statistics could be used to classify the awake and anesthetized conditions (chance level: 50%). The classification performance using the coupling to population statistic was 96.64%, that was significantly higher than using the mean activity (73.95%) or the functional connectivity (74.79%). Classification of the six experimental conditions was generally lower, but higher for *z* than for (*σ*) and FC (38% vs. 33% and 29%, chance level: 16.67%). **(D)** PCA analysis showed that *z* vectors separated the awake and anesthetized conditions along the first principal component (PC1). Each dot represents a scan. **(E)** The absolute coefficient of PC1 associated to each ROI. **(F)** During anesthesia, *z* was reduced compare to wakefulness for most of the ROIs. Changes from awake baseline, Δ*z*(*i*) = *z*(Awake) −*z*(*i*), where *z* was averaged over scans, were highly correlated for the different anesthetics (with correlation coefficients ranging from 0.85-0.93). The panel shows the comparison between Δ*z*(DPP) and Δ*z*(KET).

To examine which ROIs contributed the most to distinguish between the awake state and anesthesia based on ***z***, we performed PCA on the collection of z-scored vectors ***z***. The first principal component was sufficient to separate the awake and anesthesia conditions (**Fig. 3D**). This component had strong coefficients for brain regions located in the cingulate, parietal, intraparietal, insular cortices, and the hippocampus (**Fig. 3E**). Overall, changes in average couplings to the population with respect to awake values were similar for all anesthetics (**Fig. 3F**). We next asked how these changes affect the collective properties of brain dynamics.

### Modelling collective activity using maximum entropy models

Collective activity is ultimately described by the probability of each of the binary patterns *σ* = [*σ*_1_,…, *σ_N_*]. Estimating the distribution *P*(***σ***) over the 2^*N*^ possible binary patterns from the data is impractical with limited amount of observations, since for *N* = 82 there are more than 10^24^ possible patterns. A useful technique to estimate *P*(***σ***) relies on the maximum entropy principle. Maximum entropy models (MEMs) find *P*(***σ***) by maximizing its entropy under the constraint that some empirical statistics are preserved (see Methods). As shown above, an interesting statistic for the present study is the coupling between the state of each binary signal, *ρ_i_*, and the population activity *K*. The maximum entropy distribution that is consistent with the probability distribution *P*(*K*), the average activations 〈*σ*〉, and the linear coupling between and *K*, i.e., 〈*ρ_i_K*〉 (which relates to *z_i_*), is given by the Boltzmann distribution *P*(***σ***) = *e*^−*E*(***σ***)^/*Z*, where *E*(***σ***) represents the energy of the pattern ***σ***, given as (29):

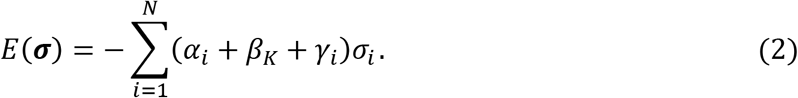

The model parameters *α_i_, β_K_*, and *γ_i_* are Lagrange multipliers associated to the constrained observables 〈*σ_i_*), *P*(*K*), and 〈*σ_i_K*〉, respectively. The normalizing constant *Z* is the partition function, given by 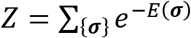, which contains information about useful statistics predicted by the model (see below). This model can be extended to include the non-linear coupling between *σ_i_* and *K*. Indeed, the maximum entropy distribution that is consistent with the joint probability distributions of and *K*, i.e., *P*(*σ_i_, K*), yields the following energy function (29):

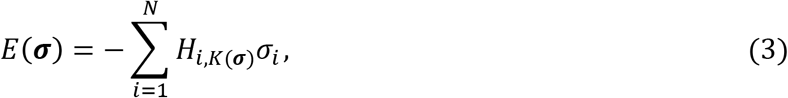

where *K*(***σ***) is the number of active ROIs in pattern ***σ*** and the parameters *H*_*i,K*(***σ***)_ represent the tendency of ROI *i* to activate when *K*(***σ***) ROIs are active. For both linear and non-linear coupling-MEMs the model parameters were inferred from the data using maximum likelihood (29). Notably, for the coupling-MEMs the partition function can be calculated directly — something that is generally not the case for most MEMs, since its calculation involves summing over all possible states.

We used these coupling-MEMs to fit the binary single-scan fMRI data for the different experimental conditions. The models accurately estimated the distribution of population activity *P*(*K*) (average Jensen-Shannon divergence *D_JS_* between the model and data distributions: *D_JS_* < 10^-6^ for both the non-linear and linear coupling-MEM; **Fig. 4A** and **Fig. S2**). Moreover, the models were able to moderately predict the covariances of the data (**Fig. 4B,C**), which were not used to constrain the models. Across the different datasets, the average correlation between the data and predicted covariances was *r* = 0.28 ± 0.03 for the linear coupling model and reached 0.40 ± 0.02 for the non-linear coupling model (see also **Fig. S2**). Furthermore, scan-classification based on parameters *γ_i_* yielded 86% and 45% correct classifications between awake and anesthetized conditions and among the six experimental conditions, respectively (**Fig. S3A,B**). Using parameters *α_i_* the classifier performance decreased to 75% and 28%, respectively (**Fig. S3B**). Thus, the learned linear coupling-MEM showed consistent variations in parameters *γ_i_* (associated to *z_i_*) across the different arousal states.

**Figure 4.**
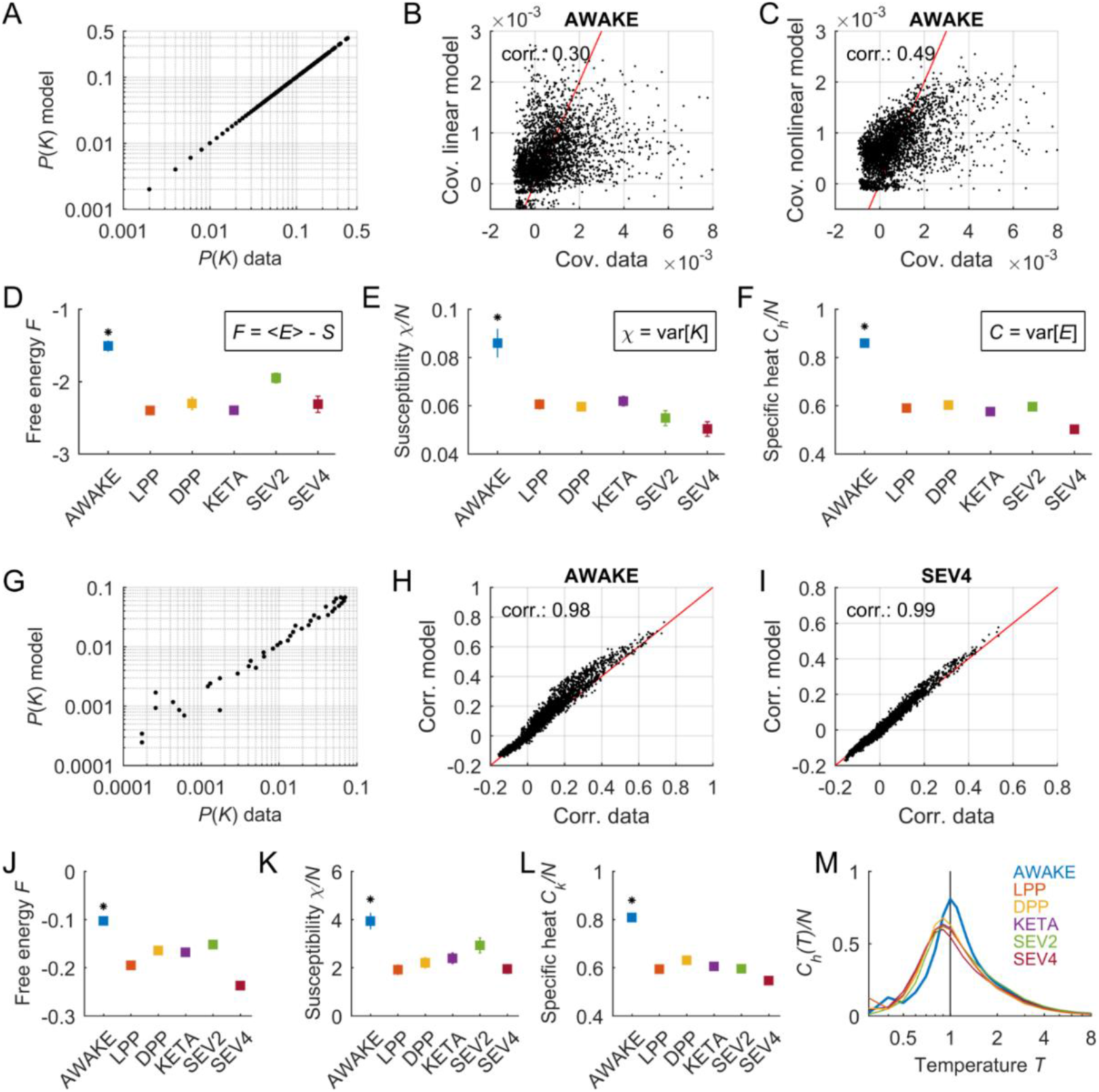
Maximum entropy models indicate higher free energy and heat capacity during wakefulness than during anesthesia. **(A)** Fitting of *P*(*K*) using the non-linear coupling-MEM. Data and predictions from all scans from the awake condition. **(B-C)** Fitting of covariances using the linear (B) and nonlinear (C) coupling-MEM for the awake condition. **(D-F)** The free energy, the susceptibility, and the heat capacity were derived using non-linear coupling-MEMs for the different conditions. Similar results were obtained using the linear model (see Fig. S4). Squares and error bars indicate means and standard deviations across scans, respectively, and the asterisks indicate significantly different values for the awake condition (p < 0.001 one-way ANOVA followed by Tukey’s post hoc analysis). **(G-L)** same as (A-F) but using pairwise-MEMs. Error bars indicate standard errors across Monte Carlo simulations of the models. Asterisks indicate significantly different values for the awake condition (p < 0.001, one-way ANOVA followed by Tukey’s post hoc analysis). **(M)** Heat capacity as a function of temperature. The peak of heat capacity for *T* = 1 indicates critical dynamics during wakefulness. The heat capacity peaked at *T* < 1 for the anesthetized conditions, indicating supercritical dynamics during anesthetized states.

### Collective activity indicated reduced free energy, susceptibility, and heat capacity under anesthesia

We can learn interesting features of collective activity using the estimated models. One important quantity is the system’s Helmholtz *free energy*, which is given by the difference between the average energy (〈*E*〉) and the entropy (*S*), i.e., *F* = 〈*E*〉 − *S*. The free energy quantifies the useful energy that is obtainable from the system. Using the Boltzmann distribution, the free energy can be directly obtained from the partition function as *F* = −ln(Z). Thus, since *Z* is tractable for the coupling-MEMs, we can directly estimate *F*. We found that the free energy was significantly higher for the awake state compared to all anesthetized conditions for both the non-linear (**Fig. 4D**) and the linear (**Fig S4A**) coupling-MEM (p < 0.001, one-way ANOVA followed by Tukey’s post hoc analysis). This result is both interesting and reasonable because it indicates that more useful energy can be extracted from the awake state than from the anesthetized state.

Two other important statistical quantities can be derived from the model, namely the *susceptibility* and the *heat capacity*. The susceptibility *χ* relates to the diversity of population states, while the heat capacity *C_h_* quantifies the diversity of accessible energy states. Specifically, the susceptibility and the heat capacity of the model are given by the variances of the population activity and the energy, respectively, i.e., *χ* = var(*K*) and *C_h_* = var(*E*). We found that *χ* and *C_h_* were significantly higher for the awake state compared to all anesthetized conditions for both linear and non-linear coupling-MEM (non-linear model: **Fig. 4E,F**, linear model: **Fig S4B,C**; p < 0.001, one-way ANOVA followed by Tukey’s post hoc analysis). This indicates that the awake system had larger population fluctuations and a larger repertoire of energy states than the system under anesthesia.

We next tested whether the same differences in these statistical quantities were found using MEM constraint by other statistics. To build the models we estimated the maximum entropy distribution *P*(***σ***) under the constraint that the activation rates (< *σ_i_* >) and the pairwise correlations (< *σ_i_σ_j_* >) are preserved. The energy of the Boltzmann distribution that is consistent with these expectation values is given by (30, 31):

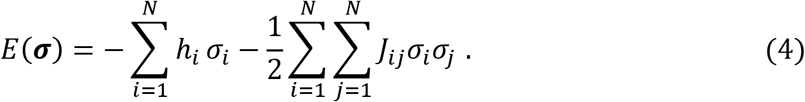

In this pairwise-MEM, the parameter *h_i_*, called intrinsic bias, represents the intrinsic tendency of ROI *i* towards activation or silence and the parameter *J_ij_* represents the effective interaction between ROIs *i* and *j*. The estimation of the model parameters **Ω** = **{*h,J*}** was achieved through a pseudo-likelihood maximization (32) (see Methods). Since this model requires the precise estimation of < *σ_i_σ_j_* >, it cannot be fitted to single-scan data and, for this reason, we used concatenated data from each experimental condition. The pairwise-MEM accurately predicted the observed correlations and, to a lower extend, it predicted the distribution of population activity *P*(*K*) (average correlation fit: *r* = 0.985 ± 0.002; average *D_JS_* = 0.006 ± 0.002; **Fig. 4G-I** and **Fig. S2**) — this is expected, since *P*(*K*) was not used to constrain the model. We found that biases and couplings parameters were changed for different states, with some parameters increasing or decreasing, and with a reduction of the variance of couplings in the anesthetized states (**Fig. S5A-D**). Moreover, coupling parameters showed a higher correlation with the anatomical connectivity (or brain connectome) in the anesthetized states than in the awake state (**Fig. S5E**).

Using this model, we calculated the collective statistical quantities for the different experimental conditions. Since in the pairwise-MEM the partition function is not tractable, we calculated *F, χ* and *C_h_* using Monte Carlo simulations (see Methods). Consistent with the above results, we found that the awake system had larger available energy (free energy, **Fig. 4J**, see also **Fig. S6**), larger population fluctuations (susceptibility, **Fig. 4K**) and larger repertoire of states (heat capacity, **Fig. 4L**) than the system under anesthesia. Thus, the different versions of the MEM used here indicate the same results concerning the statistical properties of awake and anesthetized states. Furthermore, as shown in the Appendix, the susceptibility can be viewed as a measure of the network response to a small stimulus. Consequently, we found that application of an external stimulus elicited larger and more diverse responses for the pairwise-MEM corresponding to the awake state than for the models corresponding to the anesthetized states (**Fig. S7**).

### Awake collective activity displayed critical dynamics that were shifted to a super-critical regime under anesthesia

The pairwise-MEM can be used to assess the physical state of the system. Indeed, by introducing a scaling parameter *T*, analogous to the temperature in statistical physics, one can obtain relevant features of the collective dynamics. For this, we scaled all model parameters as ***Ω → Ω**/T* and calculated the heat capacity as a function of *T*, given by *C_h_*(*T*) = var[*E*]/*T*^2^. The “temperature” *T* controls the level of disorder and its effects can be understood by examining the system’s energy levels (**Fig. S8**). Briefly, at low temperatures, interactions dominate over fluctuations making the system predominantly silent and ordered. In contrast, at high temperatures, the system is disordered and relatively uncoupled because fluctuations dominate over interactions. Both low and high temperatures lead to a low *C_h_*. However, for a specific temperature *T*_max_, order and disorder coexist in the system and *C_h_* is maximal as expected for critical dynamics (33, 34). Thus, a maximal heat capacity at *T*_max_ = 1 (corresponding to the model learned from the data) suggests that the system operated close to a critical state (whereas *T*_max_ < 1 and *T*_max_ > 1 indicates super-critical and sub-critical dynamics, respectively).

We found that the heat capacity curve was maximal for a temperature equal to 1 for the awake state, while it peaked at *T*_max_ < 1 for the anesthetized conditions (**Fig. 4M**). These results suggest that the awake state displayed critical dynamics, while dynamics under anesthesia were super-critical, which indicates that the anesthetics had a disconnection effect.

### Couplings to population relate to the sensitive parameters of the system

We next evaluated how the different parameters affected the model’s collective behavior. In general, changes in parameters can differently affect the system’s behavior, with some parameters (called “stiff” parameters) effectively modifying it, while others have little effect on it (“sloppy” parameters) (12). We used an information-theoretical approach based on the Fisher Information Matrix (FIM, noted *G)* to detect the parameters that have a strong effect on the collective activity (see Methods).

The FIM measures the change in the model log-likelihood *P*(***σ*|Ω**) with respect to changes in the model parameters **Ω**. As demonstrated in the Appendix, the FIM relates to the second derivatives of the free energy with respect to the model parameters, i.e., 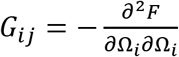. This relation provides a direct link between a macroscopic quantity (the free energy) describing the collective dynamics of the different brain states and the underlying model parameters. For the linear model, the parameters that contributed the most to the FIM were the parameters *γ_i_* (**Fig 5A**). This explains how changes in couplings to population, as observed between awake and anesthetized states, effectively change the collective state of the system, leading to the observed shift from critical to supercritical dynamics.

**Figure 5.**
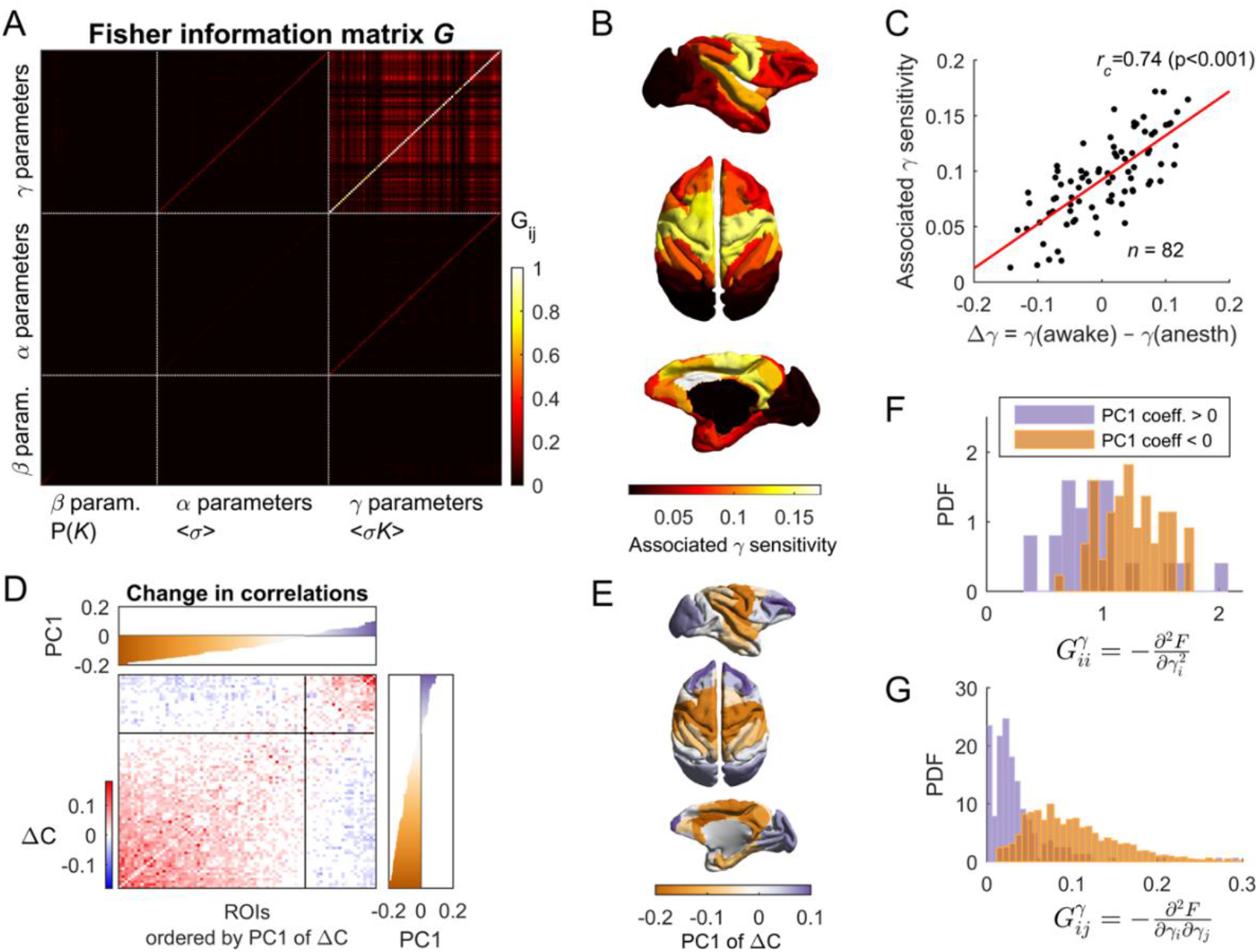
Model sensitivity to the different parameters. **(A)** Fisher information matrix (FIM) calculated using the linear coupling-MEM built using the data concatenated across scans from the awake condition. The FIM measures how much the model log-likelihood changes with respect to changes in the model parameters, i.e., **Ω** = {***α, β, γ***} for the linear coupling-MEM. **(B)** For each scan, we decomposed the FIM in eigenvectors and the mean contribution to first eigenvector of parameters *γ* was represented on the brain’s image. This represents the sensitivity of collective activity on the parameters *γ* associated to the different ROIs. **(C)** The ROIs with larger reduction of parameter *γ* in anesthetized states with respect to the awake state were those with strongest sensitivity. **(D)** Change of pairwise correlations between awake and anesthetized states: Δ*C* = *C*_awake_ − 〈*C*_anesth_〉. ROIs were ordered according to their contribution to the first eigenvector of Δ*C* (top and right insets). Two groups of ROIs were detected according to their positive or negative contribution to this eigenvector, respectively, with both groups reducing the correlation between awake and anesthesia. **(E)** First eigenvector of Δ*C* represented in the brain. **(F-G)** The two groups of ROIs had significantly different associated sensitivity (p < 0.001, Wilcoxon rank sum test), as measured by the FIM values associated to parameters *γ*.

To evaluate the importance of each of the parameters, we defined the parameter’s sensitivity as its absolute contribution to the first eigenvector of the FIM (see Methods). The regions with the largest associated sensitivity for parameter *γ* were located in the cingulate, parietal, and insular cortices (**Fig. 5B**). Those that contributed the least were visual and prefrontal cortices. Interestingly, the regions presenting larger reductions of *γ* between awake and anesthesia tended to be those with higher associated sensitivity (corr: 0.74, p < 0.001; **Fig. 5C**).

Finally, we further examined how changes in pairwise correlations between awake and anesthesia related to changes in parameters of different sensitivity. We analyzed the average difference of correlation (Δ*C*) between awake and anesthesia. Two groups of ROIs were clearly separated according to their positive or negative contribution to the first eigenvector of the matrix Δ*C*, respectively (**Fig. 5D**). Those that contributed positively were prefrontal and visual cortices, and those that contributed negatively were the cingulate, parietal, and insular cortices (**Fig. 5E**). Both groups presented a reduction of correlations under anesthesia, but prefrontal and visual cortices were related to parameters of low sensitivity (**Fig. 5F-G**). Hence, although prefrontal and visual areas changed their correlations, these changes were related to parameters that had a low impact on collective dynamics.

## Discussion

In this study, we analyzed the fMRI binary collective activity of monkeys during wakefulness and under anesthesia. We showed that the coupling between each brain region and the rest of the population provides an efficient statistic that discriminates between awake and anesthetized states. We built MEMs based on this and other statistics to derive macroscopic properties that described the different brain states, such as the free energy *F*, the susceptibility *χ*, and the heat capacity *C_h_*. All these quantities were maximized in the awake state. By studying the heat capacity curve *C_h_*(*T*) as a function of scaling parameter, controlling the disorder of the system, we showed that awake critical dynamics were shifted to supercritical ones under anesthesia. Finally, using the FIM, we showed that changes in brain state were primarily dependent on changes in the couplings to population which were associated with the sensitive parameters of the MEMs and with specific brain regions.

### Population couplings and network sensitivity

Previous research at the microcircuit level showed that neurons differ in their coupling to the population activity, with neurons that activate most often when many others are active and neurons that tend to activate more frequently when others are silent (35). Using the FIM analysis to detect sloppy and stiff parameters, it has been shown that these different types of neurons have a different impact on the network activity, different stimulus response properties, and different involvement in cortical state transitions (14). We showed that brain regions coupled differently to the rest of the whole-brain network, consistent with previous findings (28), and, furthermore, that these couplings primarily determined the collective activity (i.e., they were associated to the stiff parameters of the model) and varied across arousal brain states. Overall, we proposed that the distribution of sensitivity of brain regions and their functional role could be a general principle of neural networks at different scales.

Using principal components, we detected the combination of ROIs that contributed the most to distinguish between the awake and the anesthetized states based on their population couplings. The brain regions that changed their population coupling from the awake state to the anesthetized state were the cingulate, parietal, and insular cortices (**Fig. 3E**). Notably, the model parameters associated to the couplings of these regions were among those impacting the most the collective dynamics (**Fig. 5B-C**). Our results suggest that anesthesia modified some important local/global parameters that effectively induced a change of brain state. Our results highlight the key role of the parieto-cingulate cortex in the mechanism of anesthesia-induced loss of consciousness. Previous studies have shown that the parietal cortex (4, 9, 36) and the cingulate cortex (37) are most strongly affected by anesthetics. These cortices also present alterations in brain injury-induced unconsciousness in humans (37, 38). Moreover, consistent with our results, it has been shown that the insula plays an important role in awareness and is a potential neural correlate of consciousness (39).

Interestingly, although some brain regions, such as visual and prefrontal cortices, had different correlations between awake and anesthesia, they were associated to parameters with low impact on collective activity. This highlights the importance of studying not only the change in statistics between brain states but also their sensitivity on network dynamics. Consistent with our findings, a recent study of neuronal activity from several brain regions and in different arousal states (40) shows that parieto-basal ganglia circuits predicted the state of consciousness, while prefrontal activity failed. In addition, it has been proposed that the prefrontal cortex is mostly involved in the report of consciousness, rather than in the conscious experiences *per se* (41).

### Macroscopic *thermodynamic* quantities

Using the MEMs, we learned interesting collective properties describing the different brain states. We measured the susceptibility that quantifies the diversity of spontaneous population fluctuations. The susceptibility can be viewed as a measure of the network response to a vanishing stimulus (**Fig S7**, see also the Appendix). Thus, the higher susceptibility observed in the awake state, compared to the anesthetized states, is consistent with Transcranial Magnetic Stimulation (TMS) studies showing that stimulation elicits a more diverse and complex response in the awake state than in low-level states of consciousness, such as sleep, anesthesia, and coma (42-45). Our study predicts that the network response to a localized stimulation would covary with the population couplings and the associated parameter sensitivities of brain region.

Moreover, the models also allowed the estimation of the system’s heat capacity, a measure that quantifies the extent of the accessible dynamical repertoire. Indeed, a maximal heat capacity not only indicates that the system can display a large number of energy states, but also that these states are distinguishable (**Fig S8**). Thus, a large heat capacity indicates a large capacity to represent information in numerous separable states. The observed reduction of heat capacity in the anesthetized states is consistent with previous studies showing that the repertoire of correlation states is limited during anesthesia (11). Furthermore, by varying a scaling parameter analogous to temperature, the resulting heat capacity curves suggest that awake dynamics were critical, while anesthetized dynamics were supercritical, consistent with previous predictions (19, 20). The model used here gives an intuitive interpretation of the transition between critical to supercritical dynamics. Indeed, in the pairwise-MEM supercritical dynamics are associated with a regime in which random fluctuations dominate over interactions, which is consistent with a disconnection of *effective* couplings. It is important to note that the scale parameter *T* is only introduced to assess the state, i.e., subcritical, critical, or supercritical, of the observed system (the one given for *T* = 1, for which the pairwise-MEM fits the data). This does not mean that differences between awake and anesthetized states are due to a global reduction of interactions and biases, instead different arousal states yielded different biases and couplings (**Fig S5**) which, in combination, resulted in a change of the system’s state. This means that the anesthesia reconfigured the system and not only scaled its parameters. Consistently, we found that effective couplings correlated more with the anatomical connections for the anesthetized states than for the awake state, an effect that has been observed in empirical data (8, 11) and cannot be explained by changes in global connectivity alone (25).

Lastly, we measured the Helmholtz free energy of the estimated models. The free energy measures the useful energy that can be extracted from the system to the environment, i.e., its ability to produce work. Reasonably, the awake state led to higher free energy than the anesthetized states. Another important property of free energy is that its change with respect to the model parameters is equal to the Fisher information and, thus, it relates to the sensitivity of collective dynamics on these parameters. This result provides a direct link between the sensitivity of parameters and the change of a macroscopic quantity, the free energy, the behavior of which is known to characterize the phase transition (46). For the linear coupling-MEM, we showed that the couplings to population (*z*) were associated to the parameters that have the strongest impact on collective activity. Consistently, we found that that *z* was an efficient observable to classify the arousal states which collective dynamics were qualitatively different (in terms of criticality and supercriticality). Thus, these results give a coherent theoretical justification of the relevance of the statistic *z* to characterize the brain states and to estimate their free energy. Altogether, our findings represent a significant step in the understanding of brain states, resulting in a coherent explanation of the transition from awake to anesthesia: the phase transition between brain states is driven by those parameters that change the free energy, which are the “stiff” parameters of the systems and which relate to population couplings.

### Implications for studies on pathological low-level states of consciousness

An interesting extension of this work could be to study brain dynamics in coma using the present statistical mechanics framework. Loss of consciousness due to anesthesia or coma share common features: complexity of dynamics and neural communication are generally reduced in low-level states of consciousness (23, 47). Consequently, estimates of complexity of human brain activity have been used to assess the depth of anesthesia (48, 49) and to predict the recovery of consciousness in vegetative patients (50). Reduction of complexity is consistent with a deviation from critical dynamics when consciousness is lost. Since the coupling-MEMs can be fitted to data from single scans and, as shown here, their parameters change in different brain states, future investigation could use these models and the statistic z in the case of disorders of consciousness.

## Methods

### Animals

This study included a total of five rhesus macaques (Macaca mulatta; 4 females, 1 male, 5–8 kg; 8–12 years of age). All procedures were conducted in accordance with the European convention for animal care (86-406) and the National Institutes of Health’s Guide for the Care and Use of Laboratory Animals. Animal studies were approved by the institutional Ethical Committee (Comité d’Ethique en Expérimentation Animale, protocols #10-003 and #12-086).

### Experimental procedures

Monkeys received anesthesia either with propofol, ketamine, or sevoflurane (11). The details of the anesthesia protocols are described in the Appendix. Monkeys were scanned on a 3-T horizontal scanner (Siemens Tim Trio; TR, 2,400 ms; TE, 20 ms; and 1.5-mm^3^ voxel size; 500 brain volumes per scan session). Before each scanning session, a contrast agent monocrystalline iron oxide nanoparticle (MION) was injected into the monkey’s saphenous vein. Acquisition and preprocessing of functional images followed the standard steps described in (8) and in the Appendix. Time-series were obtained for *N* = 82 previously defined cortical regions of interest (ROIs) (CoCoMac Regional Map parcellation).

### Data binarization

fMRI time series were binarized to study the data statistics and to learn two different families of maximum entropy models (MEMs). While binarization was required to construct the MEMs, transformation of continuous fMRI signals into discrete point processes has proven to effectively capture and compress fMRI large-scale dynamics (26). Indeed, it has been shown that point process resulting from signal thresholding largely overlaps with deconvoluted fMRI signals using the hemodynamic response function and preserve the topology of the resting state networks (RSNs) [24]. We here discretized the signals as follows. For each scan, the z-scored time-series of each ROI, *x_i_*(*t*)(1 ≤ *i* ≤ *N*), was binarized by imposing a threshold *θ* = −1. Two binarization procedures were used. The first method detects the threshold crossings: the binarized activity is *σ_i_*(*t*) = 1 if *x_i_*(*t*) < *θ* and *x_i_*(*t* − 1) > *θ*, and *σ_i_*(*t*) = 0 otherwise. The second method assigns the values *1* and −1 to all time points below or above the threshold, respectively: *σ_i_*(*t*) = 1 if *x_i_*(*t*) < *θ*, and *ρ_i_*(*t*) = −1 otherwise. The first and the second procedure result in sparse and dense binary activity, respectively. We used the sparse and dense methods to construct coupling-MEMs and pairwise-MEM, respectively. This was to meet the assumptions of the model inference (see Appendix).

### *k*-means classification

We used *k*-means clustering to classify the scans based on different statistics. Let ***v***^(*i*)^ be a vector calculated from scan *i*, e.g., the vector containing all pairwise correlations among the ROIs. We used *k*-means to partition the collection of ***v***^(*l*)^ into a pre-specified number (*k*) of clusters. *k*-means minimizes the within-cluster variation, over all clusters. We used *k* = 2 to evaluate how well the scans corresponding to the awake state and those corresponding to the anesthetized states (independent of the anesthetic protocol) could be classified based on vectors ***v***^(*l*)^. To classify the six different experimental conditions, we used *k* = 6. The classification performance was given by the proportion of correctly clustered scans. We used 100 random initial conditions of the *k*-means algorithm to obtain the average classification performance and its uncertainty.

### Maximum entropy models (MEMs)

MEMs estimate the probability of all possible binary patterns, *P*(***σ***), that matches the expectation of a set of data observables. Let 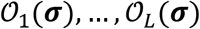 be the set of *L* data observables we seek to preserve. For example, if we were interested only on activation rates, 〈*σ_i_*〉, we would need to consider *N* observables *σ*_1_,…,*σ_N_*. Under the model distribution *P*(***σ***), the observables’ expectations are given as:

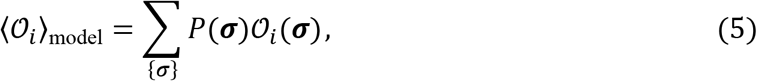

and should fit those of the data, 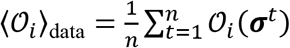, where ***σ**^t^* is the observed pattern at time *t* (1 ≤ *t* ≤ *n*). We search for the model distribution *P*(***σ***) that does less assumptions, i.e., the one that has maximal entropy *S* = ∑_{***σ***}_ *P*(***σ***)ln*P*(***σ***). Thus, the problem is equivalent to maximizing a function (the entropy) given some constraints on the expectation values of the observables, a problem that can be generally solved using Lagrange multipliers. The maximum entropy distribution has the general form:

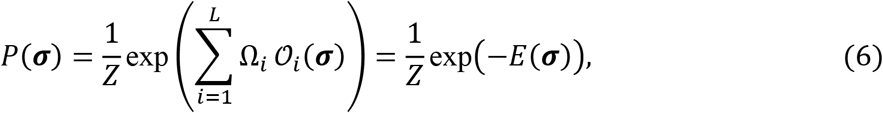

Where **Ω** = [Ω_1_,…,Ω_*L*_ are the Lagrange multipliers enforcing the constraints, 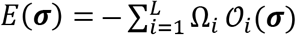 represents the energy of the pattern, and the normalizing factor *Z* = ∑_{***σ***}_ exp(−*E*(***σ***)) is the partition function (see Appendix). We estimated different MEMs built on different constrained data observables.

#### Linear coupling-MEM

First, we considered the MEM that is consistent with the probability distribution *P*(*K*), the average activations 〈*σ_i_*〉, and the linear coupling between and *K*, i.e., 〈*ρ_i_K*〉 (which relates to *z_i_*). As shown in Gardella et al. (29) the resulting energy function is given as: 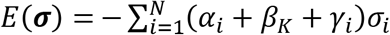. The model parameters ***α*** = [*α*_1_,…, *α_N_*], ***β*** = [*β*_0_,…, *β_N_*], and ***γ*** = [*γ*_1_,…, *γ_N_*] are Lagrange multipliers associated to the constrained observables 〈*σ_i_*〉, *P*(*K*), and 〈*σ_i_K*〉, respectively.

#### Non-linear coupling-MEM

The above model can be extended to include the non-linear coupling between and *K*. The complete coupling between *σ_i_* and *K* is provided by the joint probability distributions of *σ*_i_ and *K*, i.e., *P*(*σ_i_, K*), which is the target observable of the non-linear coupling-MEM. In this case, the energy is given as 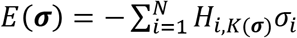 (29), where *K*(***σ***) is the number of active ROIs in pattern ***σ*** and the parameters *H*_*i,K*(***σ***)_ are associated to the constrained observables 〈*σ_i_δ_K,k_*〉, where *P*(*K = k*) = 〈*δ_K,k_*〉 and *δ_K,k_* is the Kronecker’s delta. The linear model is a special case of this model with *H*_*i,K*(***σ***)_ = *α_i_* + *β_K_* + *γ_i_*,.

#### Pairwise-MEM

The third model we considered is the one that targets the activation rates (< *σ* >) and the pairwise correlations (< *σ_i_σ_j_* >) of the data. The resulting energy function of the maximum entropy distribution is given as 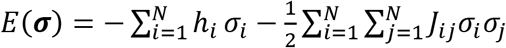 (30, 31). The model parameter *h_i_*, called intrinsic bias, represents the intrinsic tendency of neuron *i* towards activation (*σ_i_* = +1) or silence (*σ_i_* = −1) and the parameter *J_ij_* represents the effective interaction between neurons *i* and *j*.

The parameters of the coupling-MEMs and pairwise-MEMs were estimated from the data using likelihood (29) and pseudo-likelihood (32) maximization, respectively (see Appendix).

### Macroscopic quantities

The analysis of the learned MEMs provides relevant properties of the collective activity. These quantities derive from the Boltzmann distribution and they are interpretable in the framework of statistical physics. The description and calculation of these quantities are presented in the Appendix in detail. Briefly, we studied the system’s Helmholtz *free energy, susceptibility*, and *heat capacity*. The free energy *F* is given by the difference between the average energy and the entropy, i.e. *F* = 〈*E*〉 − *S* = −ln(*Z*); it quantifies the useful energy that is obtainable from the system. The susceptibility *χ* relates to the diversity of population states, i.e. *χ* = var(*K*), but, importantly, it also relates to the system’s response to intrinsic or external inputs (see Appendix and **Fig. S7**). The heat capacity *C_h_* quantifies the diversity of accessible energy states, i.e. *C_h_* = var(*E*). The heat capacity measures the size of the dynamic repertoire of the system. Furthermore, a parameter *T*, that scales all model parameters (**Ω** → **Ω**/*T*), can be introduced to study the effect of a change in the system’s disorder (“temperature”) on the repertoire of accessible energy states, i.e., the function *C_h_*(*T*) = var(*E*)/*T*^2^. This function is informative of the state of the system in terms of criticality: a maximum of the heat capacity close to *T*_max_ = 1 suggests that the observed system is likely to be close to a critical state, whereas *T*_max_ < 1 and *T*_max_ > 1 indicate super-critical and sub-critical dynamics, respectively (31, 33, 34) (see Appendix and **Fig. S8**).

### Fisher information matrix

We were interested in detecting which parameters have the strongest effect on the collective activity. For this, we studied the Fisher information matrix (FIM, noted ***G***) of the learned MEMs. The FIM represents the curvature of the log-likelihood of the model, log *P*(***σ*|Ω**), with respect to the model parameters, i.e., it quantifies the sensitivity of the model to changes in parameters. It is given as:

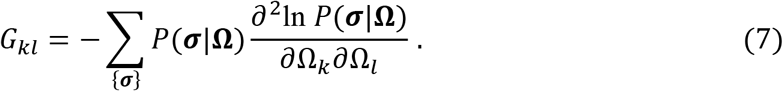

where 1 ≤ *k, l ≤ L*, where *L* is the number of parameters. As shown in Appendix, the FIM is given by the second derivatives of the free energy:

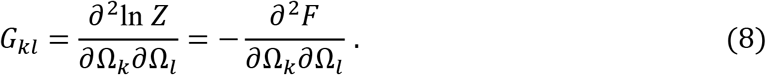

To quantify the sensitivity of the different parameters we decomposed the FIM into eigenvectors, noted ***v***_1_,…,***v**_L_*, and measured the sensitivity of a given parameter *i* by its absolute contribution to the first eigenvector, i.e. |*v*_1_(*i*)|.

### Statistical analysis

We used one-way ANOVA followed by Tukey’s post hoc analysis to compare the means of three or more distributions and Wilcoxon rank sum test to compare the medians of two distributions. We measured the dissimilarity between two distributions (i.e., data vs. model distribution) through the Jensen-Shannon divergence. Correlation matrices were analyzed using standard PCA. Statistical models (i.e., maximum entropy models) were estimated using likelihood and pseudo-likelihood maximization.

## Acknowledgements

APA, BJ, and GD received funding from the FLAG-ERA JTC (PCI2018-092891). GD acknowledges funding from the European Union’s Horizon 2020 FET Flagship Human Brain Project under Grant Agreement 785907 HBP SGA1, SGA2 and SGA3, the Spanish Ministry Research Project PSI2016-75688-P (AEI/FEDER), the Catalan Research Group Support 2017 SGR 1545, and AWAKENING (PID2019-105772GB-I00, AEI FEDER EU) funded by the Spanish Ministry of Science, Innovation and Universities (MCIU), State Research Agency (AEI) and European Regional Development Funds (FEDER). MLK is supported by the ERC Consolidator Grant: CAREGIVING (n. 615539), Center for Music in the Brain, funded by the Danish National Research Foundation (DNRF117), and Centre for Eudaimonia and Human Flourishing funded by the Pettit and Carlsberg Foundations. BJ received funding from Fondation Bettencourt-Schueller, Université Paris-Saclay (UVSQ), Fondation de France, Collége de France and from INSERM. CMS was supported by Comisión Nacional de Investigación Ciencia y Tecnología (CONICYT, currently ANID) through Programa Formacion de Capital Avanzado (PFCHA), Doctoral scholarship Becas Chile: CONICYT PFCHA/DOCTORADO BECAS CHILE/2016 - 72170507.

## Author Contributions

APA designed research, analyzed the data, studied the models, and wrote the manuscript; LU performed the experiments and curated the data; ND studied the implementation of the models; CMS curated data; MK analyzed the data and provided data visualization codes; BJ designed and supervised the experiments; GD designed and supervised research. All authors discussed the results and contributed to the editing of the manuscript.

## Supplementary Materials

## Appendix

### Anesthesia Protocols

Monkeys received anesthesia either with propofol, ketamine, or sevoflurane (11). Levels of anesthesia were defined using a clinical score and continuous electroencephalography monitoring. Under ketamine, deep propofol anesthesia and deep sevoflurane anesthesia, monkeys stopped responding to all stimuli, reaching a state of general anesthesia (11). The 3 animals scanned at a deep level of ketamine anesthesia received an intramuscular (i.m.) injection of ketamine (20 mg/kg i.m., Virbac, France), followed by a continuous intravenous infusion of ketamine (15-16 mg/kg/h i.v.) to maintain anesthesia. For propofol anesthesia, monkeys were scanned under moderate propofol sedation and deep propofol anesthesia. The awake monkeys were injected with an intravenous (i.v.) propofol bolus (5-7.5 mg/kg i.v.; Fresenius Kabi, France) to induce anesthesia, followed by target-controlled infusion (Alaris PK Syringe pump, CareFusion, CA, USA) of propofol (moderate propofol sedation: 3.7-4.0 microg/ml; deep propofol anesthesia 5.6-7,2 microg/ml). During the ketamine and moderate propofol sedation, a muscle-blocking agent was administered (cisatracrium, 0.15 mg/kg bolus i.v. followed by continuous i.v. infusion at a rate of 0.18 mg/kg/h, GlaxoSmithKline, France) to avoid artifacts related to potential movements during magnetic resonance imaging (MRI) acquisition. For sevoflurane anesthesia, monkeys were scanned under moderate and deep sevoflurane anesthesia. Monkeys received an intramuscular (i.m.) injection of ketamine (20 mg/kg i.m., Virbac, France) for induction of anesthesia, followed by sevoflurane anesthesia (moderate sevoflurane anesthesia: I/E: 2,2/2,1 vol% or deep sevoflurane anesthesia: I/E: 4,4/4,0 vol%) (Abbott, France). For all the anesthesia experiments, monkeys were intubated and ventilated. Heart rate, non-invasive blood pressure (systolic/diastolic/mean), oxygen saturation (SpO2), respiratory rate, end-tidal CO2 (EtCO2), and cutaneous temperature were monitored (Maglife, Schiller, France) and recorded online (Schiller, France).

### fMRI Data Acquisition

Monkeys were scanned on a 3-T horizontal scanner (Siemens Tim Trio; TR, 2,400 ms; TE, 20 ms; and 1.5-mm^3^ voxel size; 500 brain volumes per scan session). Before each scanning session, a contrast agent monocrystalline iron oxide nanoparticle (MION, Feraheme; AMAG Pharmaceuticals; 10 mg/kg, i.v.) was injected into the monkey’s saphenous vein (8). Preprocessing of functional images followed the standard steps described in (8), normalized to the anatomical template of the monkey MNI space (51), and band-pass filtered in the frequency range of interest (0.0025–0.05 Hz). Time-series were obtained for *N* = 82 previously defined cortical regions of interest (ROIs) (CoCoMac Regional Map parcellation). Scans that presented signs of artifacts in time-series or power spectral density were discarded. The procedure was based on the visual inspection of the time series for all the nodes, the Fourier transform of each signal. A total of 119 scans were kept for subsequent analyses, corresponding to different levels of arousal: wakefulness (*n* = 24 scans), two levels of propofol sedation (light, LPP, *n* = 21, and deep, DPP, *n* = 23), ketamine anesthesia (KETA, *n* = 22), and two types of sevoflurane anesthesia (SEV2, *n* = 18, and SEV4, *n* = 11).

### Anatomical connectivity

We used a fully weighted whole-cortex macaque structural connectivity matrix (connectome) derived by combining the information from fiber-tracing and tractography (52). The connectome is publicly available here: https://zenodo.org/record/1471588#.X44C6dAzY2x. Briefly, the tractography algorithm was optimized to best reproduce the weighted but partial-cortex tracer connectome from Markov et al. (53), before estimating whole-cortex connectome weights. The directed connectome weights between ROIs of the CoCoMac parcellation were given as the number of streamlines detected between them, divided by the total number of streamlines that were sent from the seed. Tractography-derived structural connectivity matrices were averaged across nine macaque monkeys. For details see Shen et al. (52).

### Macroscopic quantities

The analysis of the learned MEMs provides relevant properties of the collective activity. These quantities derive from the Boltzmann distribution and they are interpretable in the framework of statistical physics. Using the distribution *P*(***σ***) the mean energy and the entropy are given as:

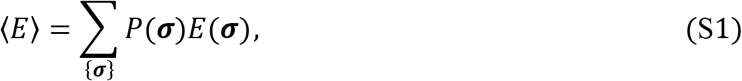

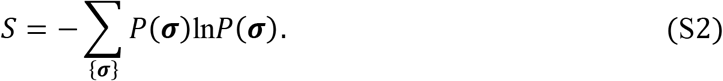

An important quantity describing the system is the Helmholtz free energy defined as: *F* = 〈*E*〉 − *S*. Using the above expressions for the mean energy and the entropy we get:

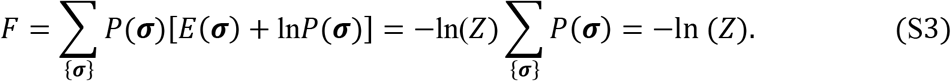

Thus, the free energy can be obtained either by calculating the mean energy and the entropy or directly by means of the partition function. These two strategies can be more or less difficult depending on the model. For the coupling-MEMs, we measured *F* for each scan through *Z*, since *Z* is tractable and the models can be fitted to single-scan data. For the pairwise-MEM, for which *Z* is not tractable, we performed Metropolis Monte Carlo simulations of the model (10^6^ steps) to estimate 〈*E*〉 and *S*. The Monte Carlo simulations were repeated ten times to estimate uncertainties on these quantities. The free energy quantifies the useful energy that can be extracted from the system, called ‘work’ in physics (54).

Another important quantity is the susceptibility *χ* that measures the fluctuations of the population activity, i.e. *χ* = var(*K*). The susceptibility can be also viewed as a measure of the population response to a perturbation applied to all units. Assuming that the perturbation *B* adds a term to the energy, i.e., 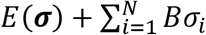, the susceptibility is given by the derivative of 〈*K*〉 with respect to *B*,i.e., *χ* = *δ*〈*K*〉/*δB* (see **Fig S7**). Here, we were interested in the so-called zero-field susceptibility obtained for *B* = 0. Using the Boltzmann distribution and noting that 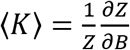 it can be shown that *δ*〈*K*〉/*δB* is equal to the variance of *K*, and thus:

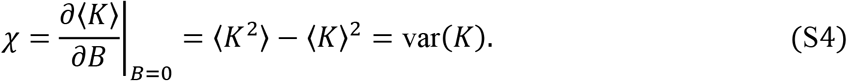

The zero-field susceptibility measures the spontaneous fluctuations of the population activity. This quantity can be measured directly in the coupling-MEMs through the estimated distribution *P*(*K*). In the pairwise-MEM, the variance of *K* can be estimated using Metropolis Monte Carlo simulations (ten simulations of 10^6^ steps). In addition, note that 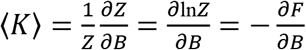 which implies the work *W* produced by changing the external stimulus from *B*_1_ to *B*_2_, i.e., 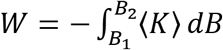, relates to the variation of the free energy Δ*F*.

A last important quantity is the heat capacity. The heat capacity *C_h_* quantifies the diversity of accessible energy states of the system, i.e., *C_h_* = var(*E*). A useful related measure is the variation of the heat capacity as a function of a scale parameter *T*, analogous to temperature in statistical physics. This parameter acts as a scaling factor for all model parameters as **Ω → Ω**/*T*. The “temperature” *T* controls the level of disorder and its effects can be understood by examining the system’s energy levels (**Fig. S8**). This creates a family of scaled models in which *T* = 1 corresponds to the MEM that was fitted to the data. The heat capacity as a function of *T* is given by *C_h_*(*T*) = var(*E*)/*T*^2^ and provides useful features of the system. Indeed, it is known that a maximum of the heat capacity close to *T* = 1 suggests that the observed system is likely to be close to a critical state, whereas *T*_max_ < 1 and *T*_max_ > 1 indicate super-critical and sub-critical dynamics, respectively (31, 33, 34). Hence, the curve *C_h_*(*T*) can be used as a tool to assess criticality. The heat capacity measures the size of the dynamic repertoire of the system. It not only provides a measure of the system’s complexity, but also assess whether the complexity is maximized and whether any reduction of complexity is due to a transition to subcritical or supercritical regimes (both regimes result in a decrease of complexity with respect to criticality, but through different mechanisms, see **Fig. S8**). For instance, if *T*_max_ ≠ 1, this means that the system can be re-scaled to increase the complexity of the model dynamics (e.g., if *T*_max_ < 1, *T* needs to decrease to reach the maximum heat capacity, indicating supercritical dynamics).

Furthermore it can be shown that the entropy *S* of the system is equal the integral of the function *C_h_*(*T*)/*T* from *T* = 0 to *T* = 1 (31, 33). This is a useful strategy to calculate the entropy that we used to compute the free energy when direct access to the partition function was not feasible (i.e., for the pairwise-MEM). In our study, we calculated *C_h_*(*T*) by estimating the variance of the energies through Monte Carlo simulations of the pairwise-MEM for different *T* (five simulations of 5.10^6^ steps for each *T*).

Finally, note that the energies are equal to the patterns’ minus log probabilities, or “surprise”, minus the free energy (a constant), i.e., *E*(***σ***) = −ln*P*(***σ***) + ln*Z*. Thus, the variance of the energy (heat capacity) measures the range of surprises of the different collective states. A large heat capacity allows the system to represent sensory events that occur with a wide range of likelihoods (energy states that are distributed, numerous, and separable) (55).

### Fisher information matrix and free energy

We were interested in detecting which parameters have the strongest effect on the collective activity. To measure how distinguishable two models, with parameters **Ω** and **Ω** + *δ***Ω**, are based on their predictions, we used the Fisher information matrix (FIM). Indeed, the Kullback-Leibler divergence between the two models can be written as:

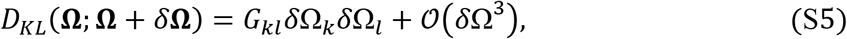

where 1 ≤ *k, l* ≤ *L*, where *L* is the number of parameters, and the matrix *G* is the FIM matrix given by:

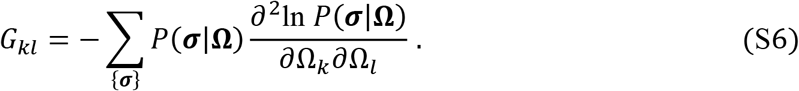

The FIM represents the curvature of the log-likelihood of the model, log *P*(***σ***|**Ω**), with respect to the model parameters, i.e., it quantifies the sensitivity of the model to changes in parameters.

Note that the FIM relates to the free energy *F* = −ln(Z). Indeed, using the Boltzmann distribution, we have:

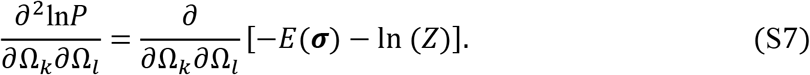

In MEMs the energy is given by the parameters (i.e., Lagrange multipliers) and the target observables as: 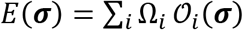. Thus, all second derivatives of the energy with respect to the parameters are zero, 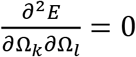, and we have:

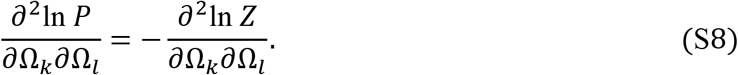

Since the right-hand term of equation 17 does not depend on ***σ***, equation 15 gives:

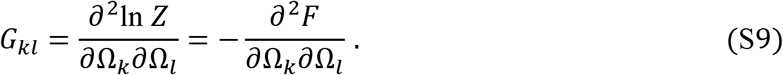

### Estimation of maximum entropy models

Linear and non-linear coupling-MEMs were estimated using the method described in (29). Briefly, Newton’s method was used to maximize the log-likelihood *LL*, by iteratively updating the parameters as:

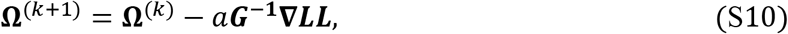

Where **Ω**^(*k*)^ is the vector of parameters at iteration *k, a* is a learning rate, ***G*** is the FIM with parameter **Ω**^(*k*)^ and **▽*LL*** is the gradient of *LL* with respect to the model parameters **Ω**^(*k*)^, i.e., 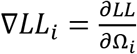. It can be shown that 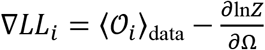. This method is feasible because, for coupling-MEM, the partition function can be analytically obtained from the model parameters and thus providing *G* and **V*LL*** see details in (29). The algorithm was stopped when the estimation errors for all constraint observables became lower than 10^-6^. We found that this algorithm correctly converged for the sparse binarization. The code for learning the coupling-MEMs is available at: https://github.com/ChrisGll/MaxEnt_Model_Population_Coupling.

The pairwise-MEM was learned using pseudo-likelihood maximization (32). This method approximates the likelihood function:

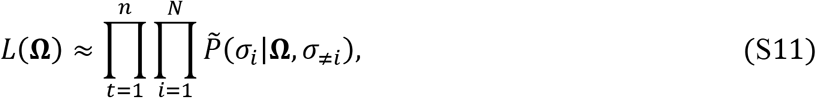

where *n* is the number of time points of the data and 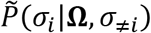 is the conditional Boltzmann distribution for a unit *σ_i_*.

This was done by updating the biases and couplings as:

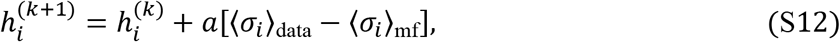

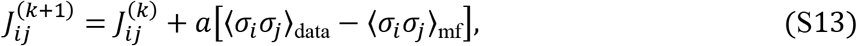

Where *k* denotes the updating iteration (up to 5.10^3^) and 〈.〉_mf_ are the expected values using the mean-field approximation:

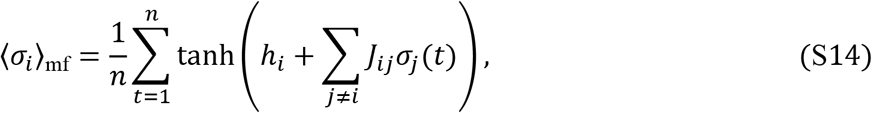

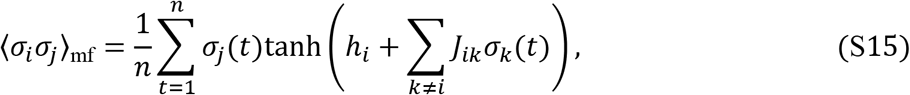

where *σ_i_*(*t*) is the activity of ROI *i* (taking values *1* or −1). The estimator of the maximum pseudo-likelihood approaches the maximum-likelihood estimator for *n* → ∞ (56). For this reason, pairwise-MEMs were fitted to concatenated data for each of the six experimental conditions. Also, since this method uses a mean-field approximation, it is accurate when the nodes of the network receive many inputs, thus we used the dense binarization scheme (see Methods). The code for learning the pairwise-MEMs is available at:

https://royalsocietypublishing.org/doi/suppl/10.1098/rsta.2016.0287.

## Supplementary Figures

**Figure S1.**
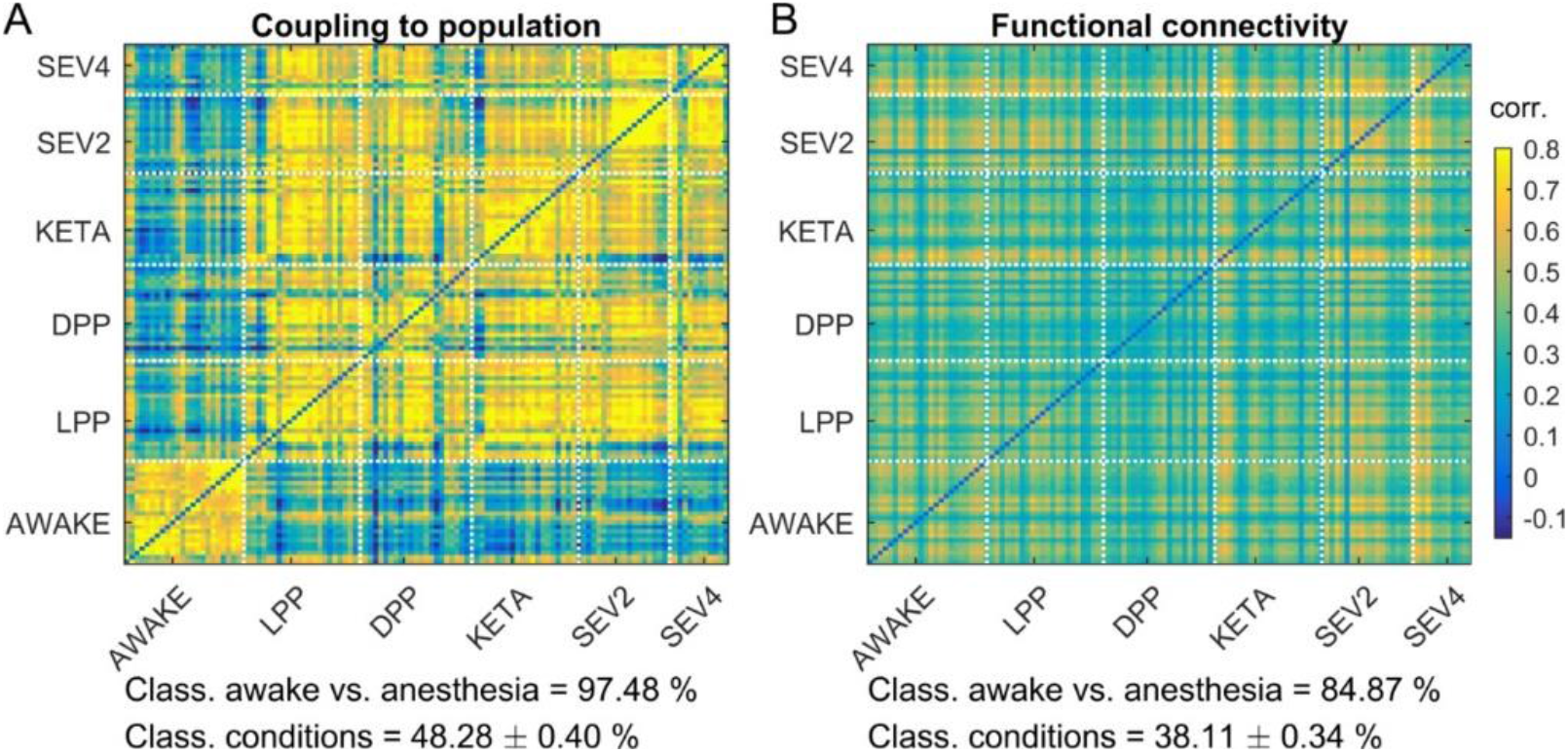
Scan-classification using continuous fMRI signals. **A-B)** Correlation matrix comparing the statistics *z* and functional correlations, calculated using continuous fMRI signals, among all scans. For example, in panel (A), the element (*k, l*) of the matrix represents the correlation between the coupling to population vector ***z*** of scans *k* and *l*. Coupling to population clearly separated awake and anesthesia data. Using *k*-means, we evaluated how well the different statistics could be used to classify the awake and anesthetized conditions (chance level: 50%). The classification performance using the coupling to population statistic was 97.48%, that was significantly higher than using the functional connectivity (84.87%). Classification of the six experimental conditions was generally lower, but higher for *z* than for functional connectivity (48% vs. 38.11%, chance level: 16.67%).

**Figure S2.**
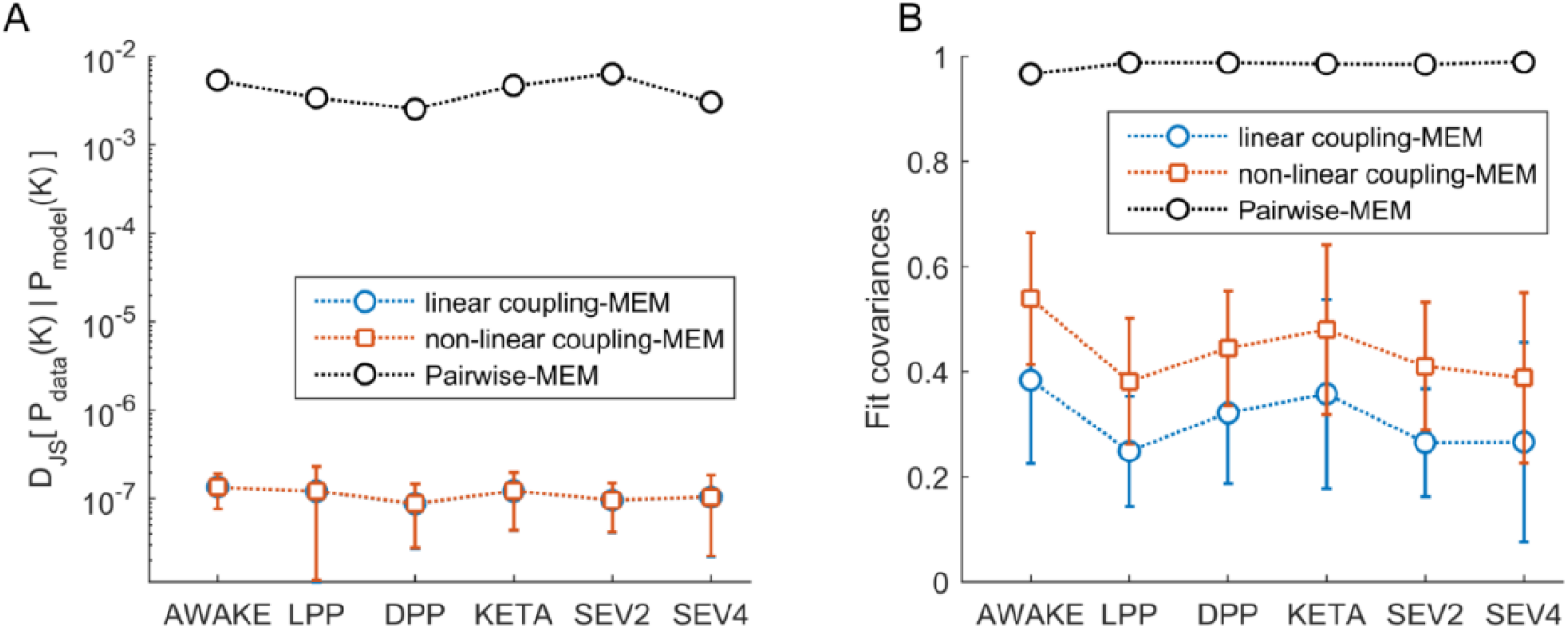
Goodness-of-fit and prediction accuracy of MEMs. We calculated the goodness-of-fit and prediction accuracy of the three types MEMs. For the linear and non-linear coupling-MEMs, the goodness-of-fit was given by the fit of the *P*(*K*) distribution, which was the target observable of the MEMs. For these models, the fit of covariances represent predictions of the models, because covariances were not used to construct them. In contrast, for the pairwise-MEM, the goodness-of-fit was given by the covariance fit (since covariances were the target of the model’s learning step) and the fit of *P*(*K*) was a model prediction. **A)** Jensen-Shannon divergence (*D_JS_*) between the model and data *P*(*K*) distributions, for the three types of MEMs (lower values of *D_JS_* indicate better approximation of *P*(*K*)). ANOVA: linear coupling-MEM: *F*_(5,118)_ = 1.3, *p* = 0.289; non-linear coupling-MEM: *F*_(5,118)_ = 1.2, *p* = 0.298. For the pairwise-MEM, ANOVA was not applicable since models were learned using concatenated data. **B)** Covariance fit for the three types of MEMs. ANOVA: linear coupling-MEM: *F*_(5,118)_ = 11.4, *p* < 0.001; non-linear coupling-MEM: *F*_(5,118)_ = 16.6, *p* < 0.001. Error bars indicate SEM.

**Figure S3.**
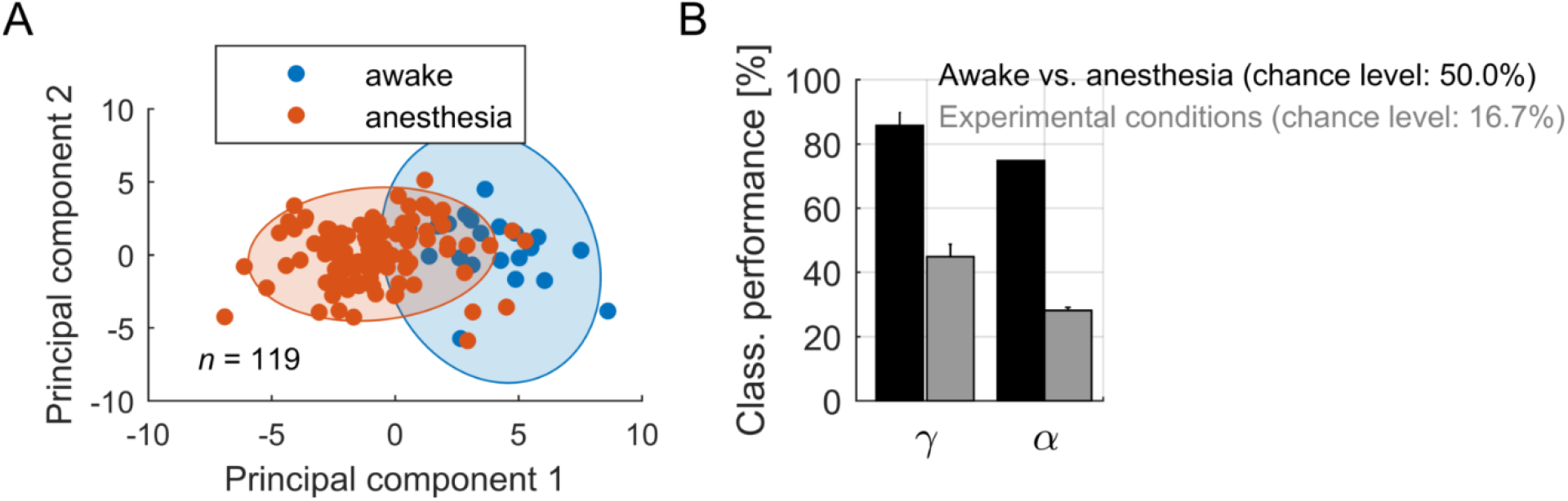
Parameters *γ* predict the state of the brain. **A)** PCA analysis showed that the parameters *γ* = [*γ*_1_,…, *γ_N_*] from the linear coupling-MEM separated the awake and anesthetized conditions along the first principal component. Each point represents a scan. **B)** Using *k*-means, we evaluated how well the different parameters *γ* and *α* could be used to classify the awake and anesthetized conditions (chance level: 50%). The classification performance based on *γ* and *α* parameters was 85.6 ± 4.4%, and 75.8 ± 0.0%, respectively. Classification of the six experimental conditions was 45.0 ± 4.0%, and 27.9 ± 0.9% based on *γ* and *α* parameters, respectively.

**Figure S4.**
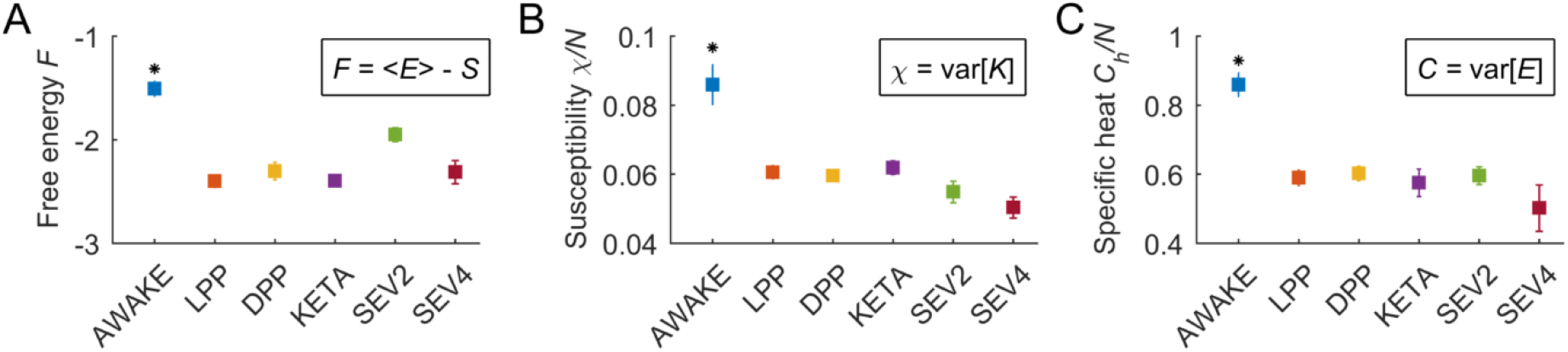
Macroscopic quantities using the linear coupling-MEM. **A)** Free energy *F* for the different conditions, *F* = 〈*E*〉 − *S* = −ln *Z*. **B)** Susceptibility *χ* for the different conditions, *χ* = var(*K*). **C)** Heat capacity *C* for the different conditions, *C_h_* = var(*E*). In panels (A) to (B), we the linear model was used. Squares and error bars indicate means and standard deviations across scans, respectively, and the asterisks indicate significantly different values for the awake condition (p < 0.001 one-way ANOVA followed by Tukey’s post hoc analysis).

**Figure S5.**
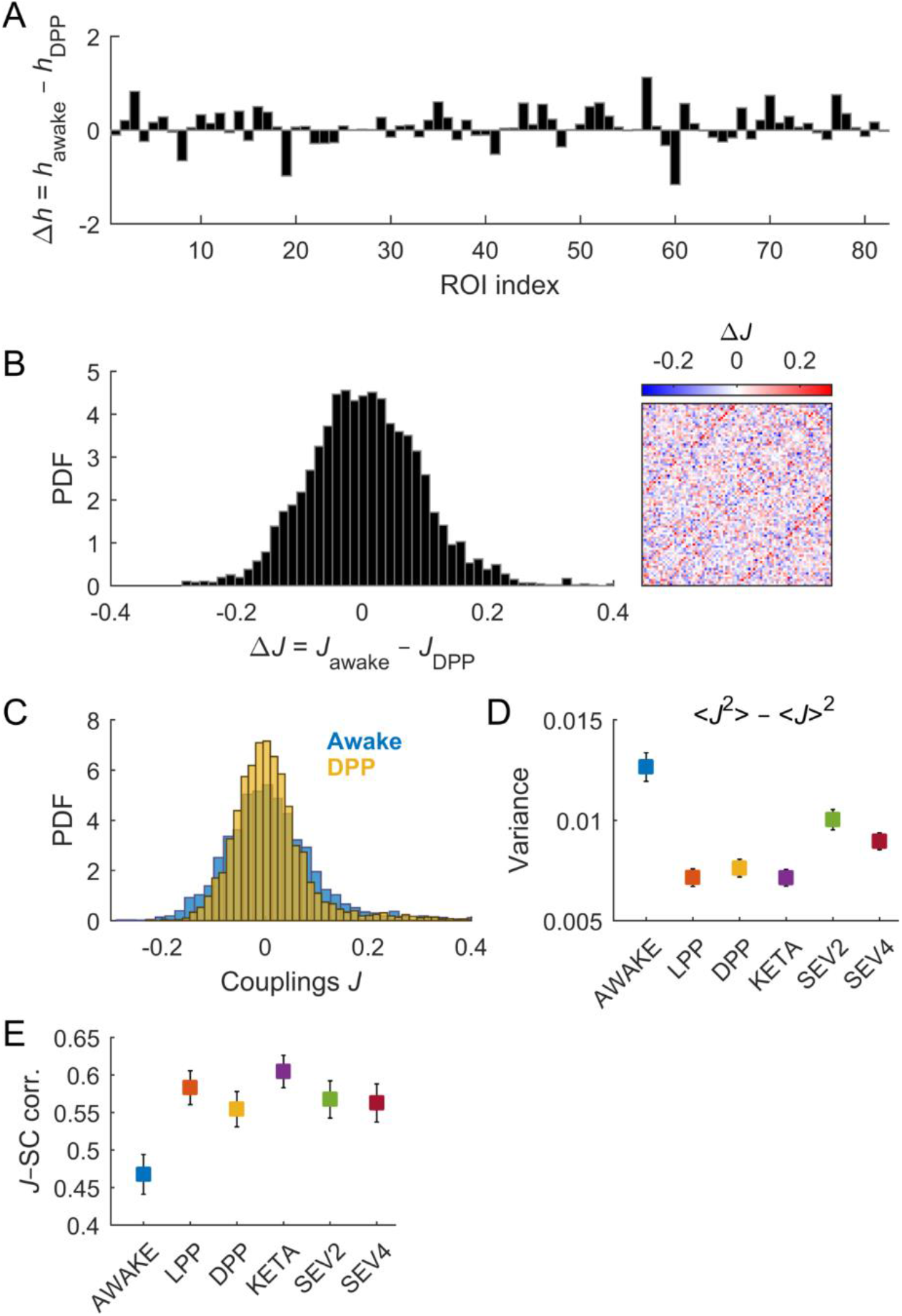
Change of the pairwise-MEM parameters *h* and *J* across brain states. **A)** Change in bias model parameters (*h*) between the awake and deep propofol (DPP) conditions for each ROI. **B)** Distribution of the change in couplings (*J*) between the awake and deep propofol (DPP) conditions for all pairs of ROIs. Inset: the matrix represents the change of the couplings between awake and DPP, i.e., 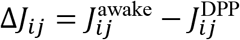. **C)** Distribution of couplings *J* in the awake and DPP conditions. **D)** Variance of couplings *J* for each experimental condition. Error bars indicate bootstrap uncertainties (500 repetitions). **E)** Pearson correlation between the coupling matrix *J* and the structural connectivity, for each experimental condition. The error bars indicate the correlation coefficient’s 95% confidence interval.

**Figure S6.**
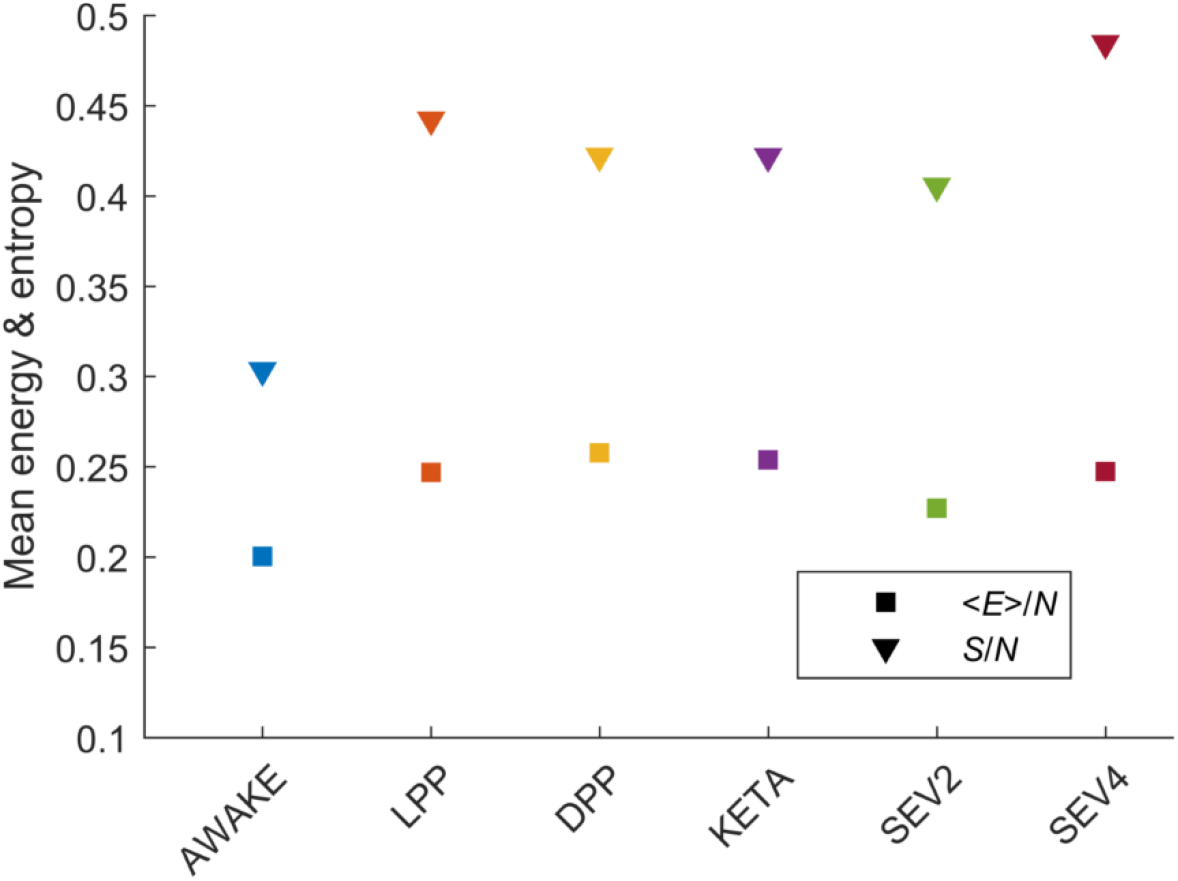
Mean energy and entropy for the pairwise-MEM. The mean energy 〈*E*〉 and the entropy *S* of the learned pairwise-MEMs were calculated using Monte Carlo simulations (10^6^ steps). To calculate the entropy, we used the heat capacity as a function of the scaled parameter *T*, analogous to temperature (see Methods).

**Figure S7.**
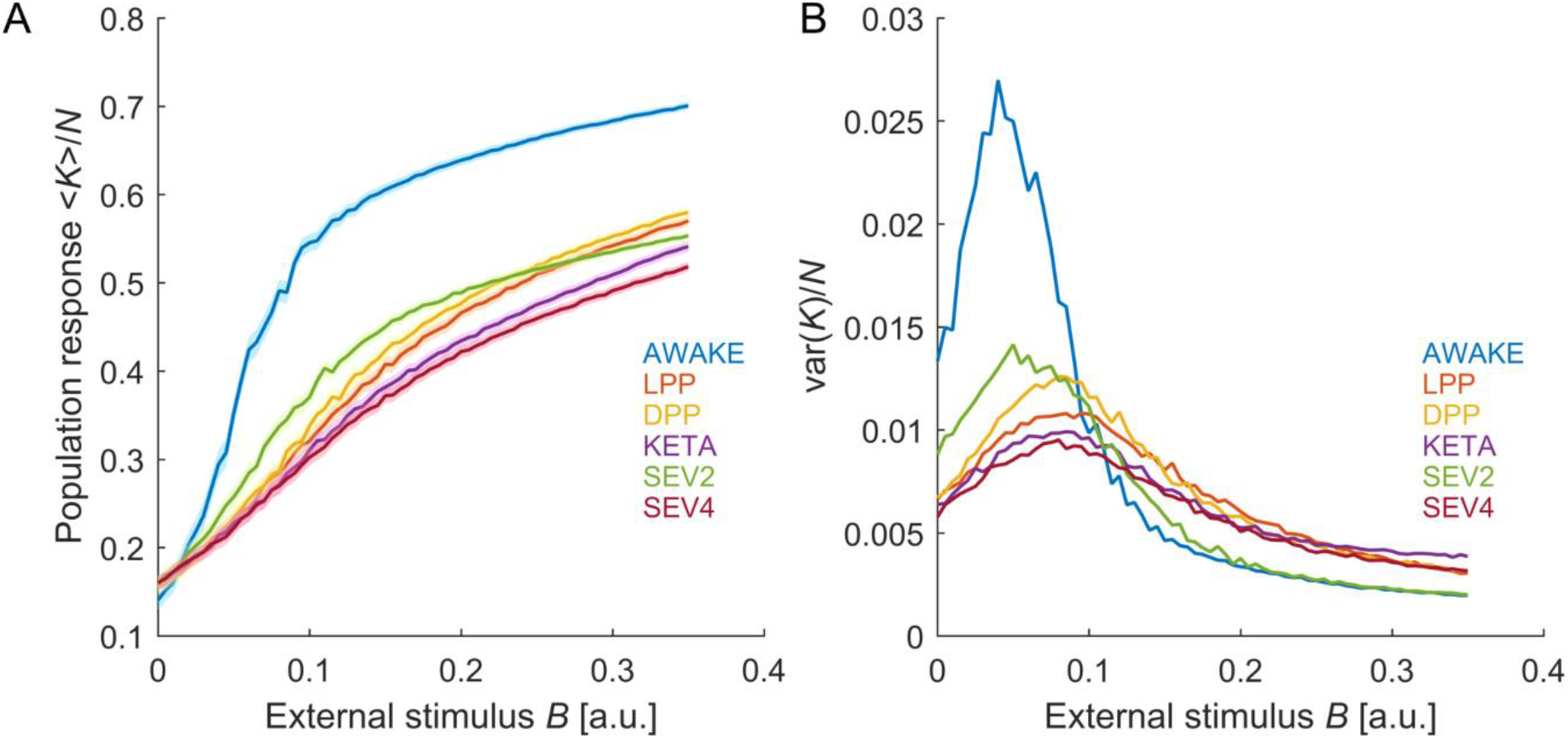
Population response to an external stimulus predicted by the pairwise-MEMs. **A)** An external stimulus was applied to the learned pairwise-MEMs. The stimulus *B* added a term to the energy as: 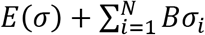. We performed Monte Carlo simulations (100 trials of 5.10^4^ steps) for different values of *B* to examine the mean population response 〈*K*〉 as a function of the external stimulus. In this case, the work *W* produced by changing the external stimulus from *B*_1_ to *B*_2_, i.e., 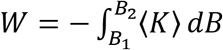, relates to the variation of the free energy Δ*F*. **B)** Variance of the population activity. In these simulations, inactive ROIs (*σ_i_* = −1) were set to 0, so that *K* represents the number of active ROIs.

**Figure S8.**
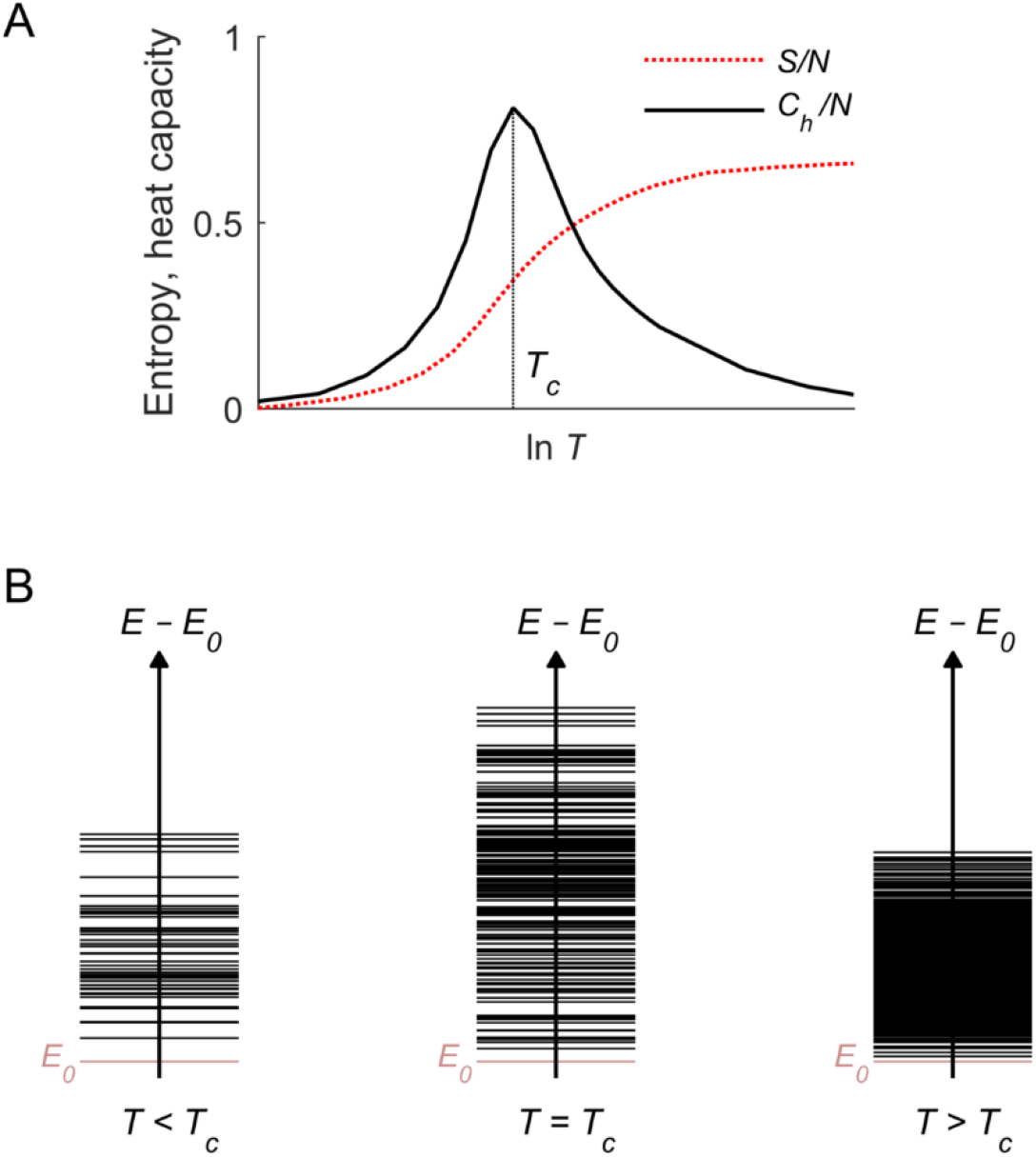
Heat capacity and energy levels. **A)** Heat capacity *C_h_* and entropy *S* of the pairwise-MEM, as a function of the scalling parameter *T*, analogous to temperature in statistical physics. The temperature *T* controls the disorder of the system (i.e., the entropy *S*(*T*) increases as a function of *T*). If the model learned from the data (i.e., *T* = 1) maximized the heat capacity, then its dynamics are critical. If the temperature that maximizes the heat capacity is larger than 1, the dynamics of the learn model are subcritical (the temperature must increase to reach the maximum). Finally, if the temperature that maximizes the heat capacity is lower than 1, the dynamics of the learn model are supercritical (the temperature must decrease to reach the maximum). **B)** Observed energies *E* for subcritical, critical, and supercritical dynamics. *E*_0_ correspond to the energy of the state for which all units are silent. For subcritical dynamics, the few visited energy levels are sparsely distributed, and the variance of energies (i.e., heat capacity) is relatively low. For supercritical dynamics, many energy levels are densely distributed, and their variance of energy is also relatively low. For critical dynamics, the energy levels are numerous and separable, leading to a maximal variance of energies.

